# SpatioCell: Deep Integration of Histology and Spatial Transcriptomics for Profiling the Cellular Microenvironment at Single-Cell Level

**DOI:** 10.1101/2025.04.07.647590

**Authors:** Naiqiao Hou, Yue Yu, Zhaorun Wu, Yanqi Zhang, Jiayun Wu, Zhixing Zhong, Fangfang Xu, Zeyu Wang, Chaoyong Yang, Weihong Tan, Jia Song

## Abstract

Spatial transcriptomics (ST) is a powerful assay to capture gene expression in tissue context. However, due to the limitation of resolution, most existing ST datasets remain at multicellular resolution which hinders comprehensive understanding of the spatial organization. We propose SpatioCell, a computational algorithm to automatically extract both cell type and expression information at single-cell resolution from ST data, through a morpho-transcriptomic spatial reconstruction framework solved via dynamic programming, integrating morphological and transcriptomic information. This framework enables deterministic single-cell spatial reconstruction, assigning cell identities to precise locations rather than inferring spot-level cell-type composition and expression, and further uncovers overlooked microenvironment features while correcting deconvolution errors. Using the ST data from triple negative breast cancer as an example, SpatioCell reveals the significance of the distance between cancer-associated fibroblast (CAF) and tumor or immune cells for disease progression. The establishment of SpatioCell will broaden the biomedical applications of ST and facilitate investigations of single-cell spatial organization.

## Background

Deciphering the cellular composition of tissues is crucial for understanding how spatial organization dictates cell interactions, functional specialization, and microenvironmental dynamics. These structural complexities shape physiological processes and influence disease progression, immune responses, and therapeutic outcomes^1, 2^.

Spatial transcriptomics (ST) has emerged as a powerful tool for profiling gene expression within the spatial context of tissues, enabling the characterization of cell compositions and molecular landscapes in situ^3, 4^. Existing ST technologies can be broadly categorized into image-based and sequencing-based approaches^5^. Image-based methods, such as MERFISH^6^ and seqFISH+^7^, achieve subcellular resolution, but they are limited to predefined gene sets. In contrast, sequencing-based methods, including 10x Visium^8^, Decoder-seq^9^, and Stereo-seq^10^, provide whole-transcriptome coverage with higher throughput and simpler protocols, making them more widely adopted. However, their resolution remains constrained, with platforms like Visium featuring spot sizes around 55 μm, often capturing multiple cells within each spatial unit (hereinafter termed as a spot). As most current ST datasets are derived from these sequencing-based platforms, the growing volume of data, while offering valuable insights into tissue organization, still largely remains at multicellular resolution, hindering single-cell level analysis ^11^.

To address these resolution constraints, computational methods have been developed. Deconvolution algorithms leverage single-cell RNA sequencing (scRNA-seq) references to infer cellular composition within each spatial spot^12, 13^. However, these approaches estimate only cell-type proportions rather than achieving actual single-cell resolution, which limits precise spatial mapping and hinders comprehensive characterization of microenvironmental heterogeneity, especially in terms of the fine-scale spatial distribution, intercellular distances, and cell interactions.

Notably, ST data is often accompanied by histology images, mostly through hematoxylin and eosin (H&E) staining, which provide additional spatial context that could enhance resolution. Histology image analysis, a key focus in computational pathology, has led to the development of several cell/nuclear segmentation models^14, 15^. Current studies (such as SpatialScope^16^ and Spotiphy^17^) have integrated these segmentation models with ST data analysis. However, these approaches are mainly limited to either counting the number of nuclei within each spot or using histology-derived information to heuristically distribute spot-level gene expression in pursuit of super-resolution profiles, but they still cannot precisely assign cell types to spatial positions. This limitation largely arises because few models can accurately perform both nuclear segmentation and cellular morphological analysis on H&E-stained images. Moreover, no effective annotation methods are available to integrate image-derived morphological features with sequencing information.

To address these challenges, we developed SpatioCell, a morpho-transcriptomic spatial reconstruction framework that produces high-resolution, single-cell–resolved maps of both cell types and their gene expression. SpatioCell re-engineers the Segment Anything Model (SAM) ^18^, an instance segmentation foundation model, by redesigning its prompting^19^ and decoding mechanisms to achieve robust and accurate nuclear segmentation and morphological analysis in complex tissues. It further models the cell-type assignment, based on histology-derived cell-identity features and deconvolution predictions, as a constrained combinatorial optimization problem, and employs a dynamic programming (DP) algorithm to achieve precise single-cell annotation. In extensive evaluations on six public datasets comprising thousands of tissue-stained images or ST slides, SpatioCell outperformed state-of-the-art methods in nuclear segmentation and classification. Applied to tumor samples with multicellular-resolution ST (55–100 μm), it corrected deconvolution errors and reconstructed single-cell maps of cell type and expression, revealing tumor microenvironment (TME) features undetectable by existing approaches. In triple-negative breast cancer (TNBC), SpatioCell identified cancer-associated fibroblast (CAF) subtypes with distinct spatial proximities to tumor and immune cells. By integrating intercellular distance, gene expression, and clinical outcome data, it further revealed functionally and prognostically relevant CAF states shaped by the local microenvironment. These findings demonstrate the utility of SpatioCell as a powerful framework for decoding microenvironment-driven cellular heterogeneity, and further suggest its potential to advance the biomedical applications of ST by enabling accurate analysis of cell–cell distances, interactions, functions, and their implications in disease.

## Results

### Overview of SpatioCell

To achieve single-cell–level cell annotation in multicellular-resolution ST data, existing algorithms use histopathological images to locate and count cells within each spot but often overlook the rich morphological cues, leaving precise single-cell mapping beyond the reach of current methods. Here we developed SpatioCell, by integrating morphological information from histopathological images, it enables single-cell–resolved cell-type assignment and elevating gene expression profiles to the single-cell level. SpatioCell comprises two key modules: (1) a specialized spatio-morphological learner for histopathological image processing, which performs nuclear segmentation and detailed morphological feature extraction, and (2) a morpho-transcriptomic spatial reconstruction module for cell annotation that integrates image-derived morphological features with transcriptomic data to achieve high-precision, single-cell–level annotation (**Fig. 1A**).

**Figure 1.**
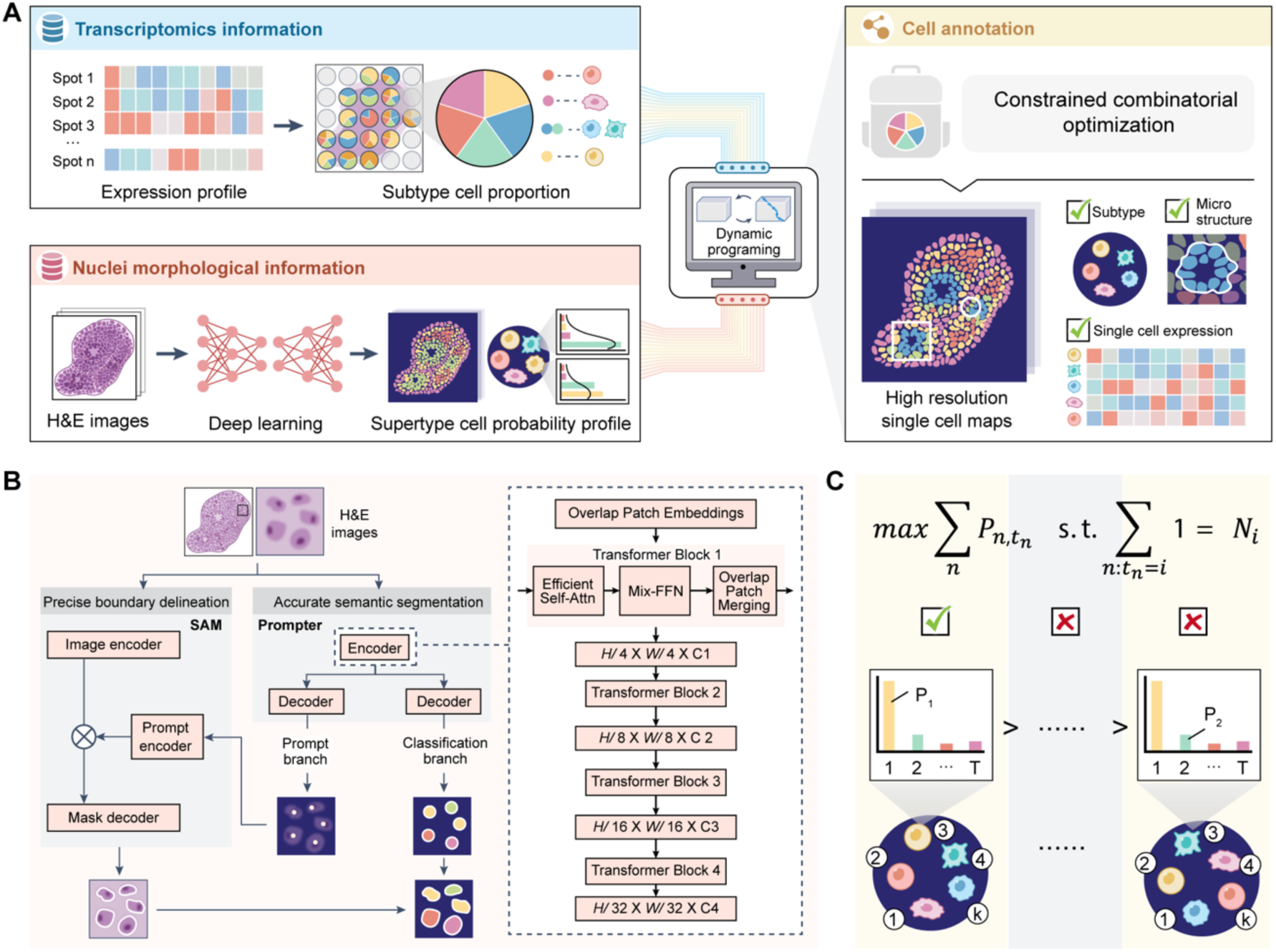
Overview of SpatioCell. **(A)**, SpatioCell computational framework. SpatioCell consists of two main modules: (1) a specialized spatio-morphological learner, which is a deep learning module for histopathological image processing that performs nuclear segmentation and analysis through detailed morphological feature extraction, and (2) a morpho-transcriptomic spatial reconstruction module which is a DP module for cell annotation and gene expression mapping that integrates supertype and subtype annotations derived from morphological and transcriptomic data to achieve high precision and resolution. Module 1 generates a probabilistic profile for each mask based on its morphological features, allowing SpatioCell to model the integration as a knapsack problem. **(B)** Deep learning architecture of module 1. Left: The framework includes (1) a fine-tuned SAM-based segmentation model and (2) an adaptive prompting mechanism, which allows SAM to receive accurate positive prompts for automated nuclear segmentation. The final outputs include cell locations, masks, and cell identity (supertype) probability profiles derived from morphological cues. Right: Detailed architecture of the Prompter. The Prompter’s encoder uses overlapping patch embedding to extract hierarchical features. This includes (1) Efficient Self-Attention for long-range dependencies, (2) Mix-FFN for feature transformation, and (3) Overlap Patch Merging for spatial dimension reduction. The output feature maps enable nuclear center detection and semantic segmentation. **(C),** Schematic illustration of the optimization problem of module 2. The goal of cell type assignment is to maximize the total type probability within each spot, where *P*_*n,t,n*_ denotes the probability that cell *n* is classified as type *t*_*n*_, subject to the constraint that the number of cells assigned to each type i equals a predefined value *N*_*i*_. The left panel shows an optimal assignment maximizing the summed probabilities, while the right panels illustrate suboptimal assignments under the same constraints.

Conventional SAM, while powerful in natural image segmentation^18^, struggles to generalize to H&E-stained pathology images due to blurred nuclear boundaries, heterogeneous cell morphologies, and densely packed tissue architectures^20^. To address these limitations, we propose a spatio-morphological learner within SpatioCell, a pathology-tailored segmentation framework that systematically re-engineers SAM’s prompting and decoding mechanisms. Unlike prior adaptations that only fine-tune SAM, SpatioCell introduces three key innovations. First, it leverages nuclear mask– derived prompts during fine-tuning, aligning the foundation model with histopathology-specific object scales and boundaries. Second, we design an automatic Prompter, built upon a multi-scale transformer encoder^19^, that simultaneously encodes global tissue context and fine-grained nuclear details to generate precise positive prompts without manual intervention. Third, we augment the Prompter with a semantic classification head, enabling joint modeling of nuclear segmentation and cell-type identification within a unified architecture (**Fig. 1B**). Through this design, SpatioCell outputs both (i) accurate nuclear masks and (ii) cell identity (supertype) probability profiles. This reconfiguration of SAM into a specialized spatio-morphological learner establishes a new paradigm for foundation models in computational pathology.

While H&E images provide valuable insights into cell localization and coarse cell supertypes, they lack the resolution to resolve fine-grained subtypes and are often noisy, necessitating integration with ST. Conventional ST analysis relies on deconvolution algorithms to estimate the aggregate composition of cell subtypes within each spot, but these approaches operate only at spot resolution. SpatioCell introduces a novel optimization-based framework to bridge this gap by coupling morphology-derived probability profiles with deconvolution-derived compositional constraints. Specifically, the spatio-morphological learner outputs a probability distribution across cell types for each nucleus, based on which we reformulate the cell identity assignment as a constrained combinatorial optimization problem (i.e. a knapsack problem) (**Fig. 1C**): the objective is to maximize the global likelihood of type assignments under limited capacity defined by deconvolution-inferred cell-type proportions. This formulation enables a dynamic programming (DP) solver to jointly optimize assignments across all cells in a spot, ensuring consistency with global transcriptomic priors while leveraging fine-grained morphological cues. Lastly, by mapping the cell identities with DP and modeling the spot-level counts as a multinomial distribution of each gene’s reference profiles weighted by the cell-type proportions^17^, we further resolve the inferred gene expression for each cell at specific spatial location, thereby generating a high-precision and high-resolution atlas of cell identity and gene expression. In contrast to prior mapping approaches, SpatioCell’s optimization-driven integration achieves both fidelity to transcriptomic constraints and resolution at the single-cell level, establishing a new paradigm for cross-modal reconstruction in ST.

Above all, SpatioCell introduces a framework that integrates histological and transcriptomic data to enhance ST resolution to the single-cell level. It refines nuclear segmentation with a specialized spatio-morphological learner and assigns cell identities and gene expression using a constrained optimization strategy, ensuring precise and robust annotation while achieving high-resolution spatial mapping of transcriptomic profiles.

### SpatioCell Achieves Superior Nuclear Segmentation on H&E Images from ST Data

To assess the effectiveness of SpatioCell’s image processing module, we benchmarked it against five widely used nuclear segmentation and classification algorithms, including Hover-Net^21^, Cellpose3^22^, StarDist^23^, Mask R-CNN^24^, and CellProfiler^25^. Evaluation metrics included the Aggregated Jaccard Index (AJI), Panoptic Quality (PQ) and its components (Detection Quality, DQ and Segmentation Quality, SQ), as well as the detection ratio for instance segmentation. For nuclear type classification, mean PQ (mPQ) per class and a classification-aware F-score F_C_, proposed by Graham et al., were used^21^. These metrics provided a comprehensive assessment of both instance segmentation accuracy and semantic segmentation performance (see **SI Note 1**. Evaluation Metrics for details).

These models were first tested on four public H&E-stained histopathology datasets for the instance segmentation task (see **SI Note 1**. Dataset for details). SpatioCell achieved the best performance across most evaluation metrics in CPM17, Kumar and CoNSeP datasets, including in challenging cases with densely packed and overlapping nuclei, further demonstrating its robustness and reliability (**Fig. 2A**, **Fig. S1**, **Table S1** to **3**). In PanNuke, one of the most complex benchmarks with over 200,000 annotated nuclei across 19 tissue types^26^, SpatioCell consistently outperformed all other models in both nuclear detection and segmentation metrics across three cross-validation folds (**Fig. 2B**, **Table S4**). Furthermore, SpatioCell both successfully detected nuclei with weaker staining and accurately excluded unstained mature red blood cells (**Fig. 2C**), demonstrating its ability to remove background artifacts and accurately segment true nuclei.

**Fig. 2.**
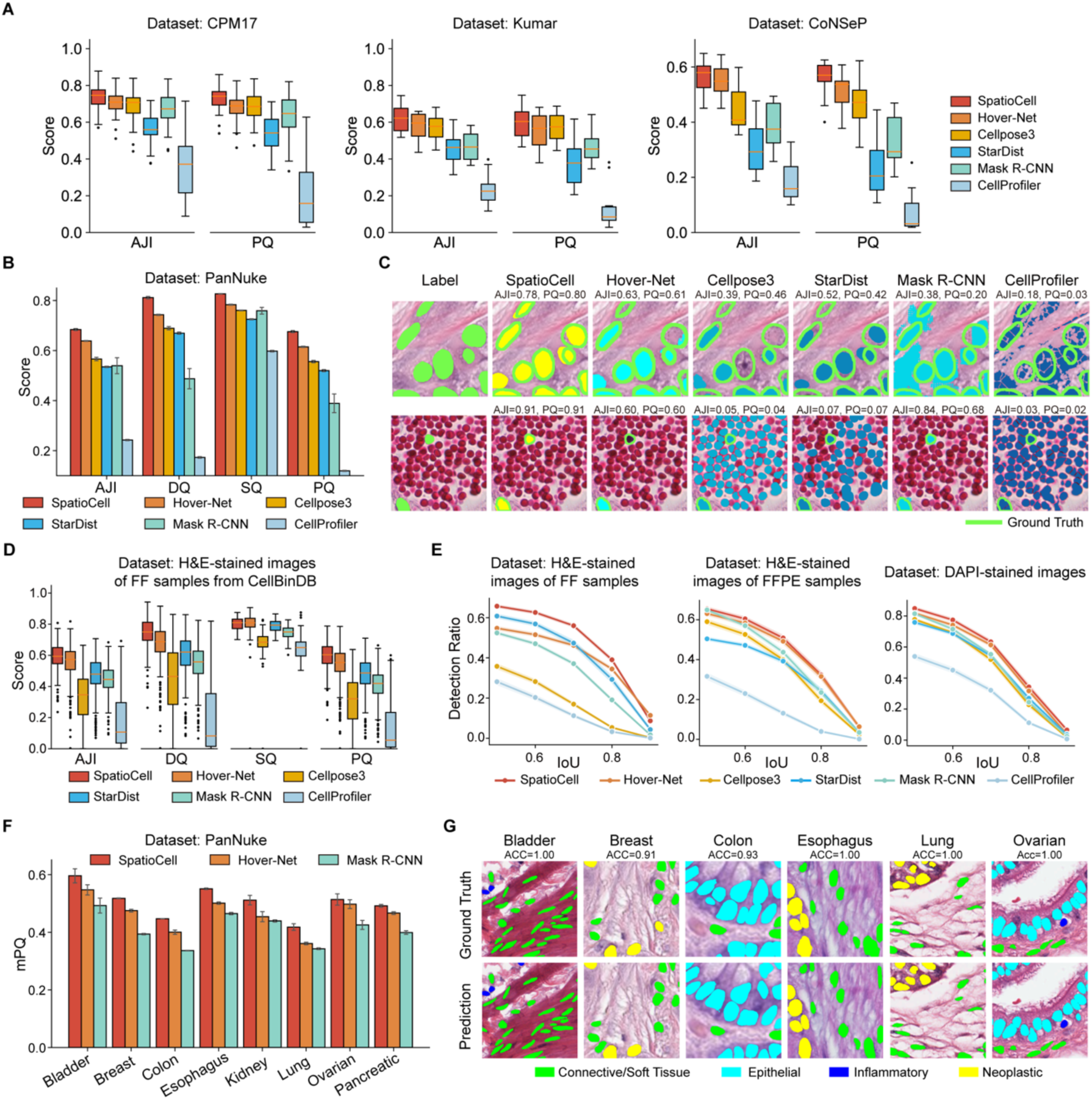
Performance evaluation of SpatioCell in nuclear segmentation and analysis. **(A)**, Segmentation performance comparison of SpatioCell with other 5 methods on CPM17 (left), Kumar (middle) and CoNSeP (right) datasets. Each data point represents a metric calculated from a sample image in the test datasets. **(B)**, Segmentation performance of various methods on the PanNuke dataset via three-fold cross-validation. Error bars represent standard error. **(C)**, Visual examples of nuclear segmentation on the PanNuke dataset, with AJI and PQ scores annotated. **(D)**, Nuclear segmentation performance of different methods on H&E-stained images of FF samples from the CellBinDB dataset. **(E)**, Detection ratio of various methods on the CellBinDB dataset under different IoU thresholds. **(F)**, Distribution of mPQ scores of SpatioCell, Hover-Net, and Mask R-CNN on the PanNuke dataset through three-fold cross-validation. Nuclear type classification for SpatioCell is based on the maximum probability in the classification probability profile. **(G)**, Visual examples of nuclear classification by SpatioCell on the PanNuke dataset.

To further assess its generalizability to ST datasets, we conducted benchmark experiments on the CellBinDB dataset, which includes H&E-stained images from both formalin-fixed, paraffin-embedded(FFPE) and fresh frozen(FF) preparations, covering both normal and diseased tissues from human and mouse samples^27^. SpatioCell consistently achieved the highest scores across all evaluation metrics, demonstrating its robustness in diverse histopathological images, particularly excelling in the segmentation of FF H&E images, where nuclei are more faint and morphologically variable (**Fig. 2D** and **E**, **Fig. S2A** to **C**). The accuracy of nuclear detection is critical for SpatioCell’s downstream cell annotation and quantitative analysis across multiple IoU thresholds confirmed SpatioCell’s superior nuclear detection capabilities, especially on small and faint nuclei (**Fig. 2E**, **Fig. S2B**). Furthermore, we tested SpatioCell on DAPI-stained images from the dataset, a common imaging type in ST, and found it generalizes well across different staining methods (**Fig. 2E**, **Fig. S2A** and **D**).

To incorporate additional morphological information and enhance cell annotation, SpatioCell was further developed to include a nuclear classification module, providing essential input for the subsequent cell annotation step. To evaluate its effectiveness, we compared SpatioCell with Hover-Net and Mask R-CNN, two widely used semantic segmentation models, across three benchmark datasets: PanNuke, MoNuSAC, and CoNSeP. SpatioCell outperformed both methods across all three datasets (see results of CoNSeP and MoNuSAC in **Table S5-6**). Specifically, in three-fold cross-validation on PanNuke, SpatioCell achieved the highest F_C_ compared to the other two methods (**Table S7**). We also examined the mPQ per tissue type, and SpatioCell outperformed both methods in18 out of 19 tissues (**Fig. 2F**, **Fig. S3**). As shown in **Fig. 2G**, SpatioCell demonstrates robust classification capability across diverse histopathological images.

### High-Precision Single-Cell Annotation and Gene Expression Mapping in Spatial Transcriptomics

To overcome the limitation of multicellular resolution, the cell annotation module of SpatioCell combines H&E-derived and expression-derived information, enabling the construction of high-resolution single-cell spatial maps and improving the accuracy of cell-type assignment in STs. (**Fig. 3A Methods**). To comprehensively evaluate SpatioCell’s annotation capability in ST data, we simulated multicellular-resolution ST datasets based on single-cell resolution ST (10x Xenium) from breast cancer (BRCA), ovarian cancer (OVCA), pancreatic cancer (PDAC) and lung cancer (LUCA). We compared annotation results from SpatioCell with two widely used approaches: an H&E image-based classification and a deconvolution-based method that randomly assigns identities to the segmented cells within each spot based on cell type proportions, representing advanced deconvolution-based strategies such as those used in Spotiphy^17^, as detailed in **Methods**. This comparison allows us to assess whether the deep integration of H&E-derived and expression-derived information in SpatioCell outperforms the use of either data type alone.

**Figure 3.**
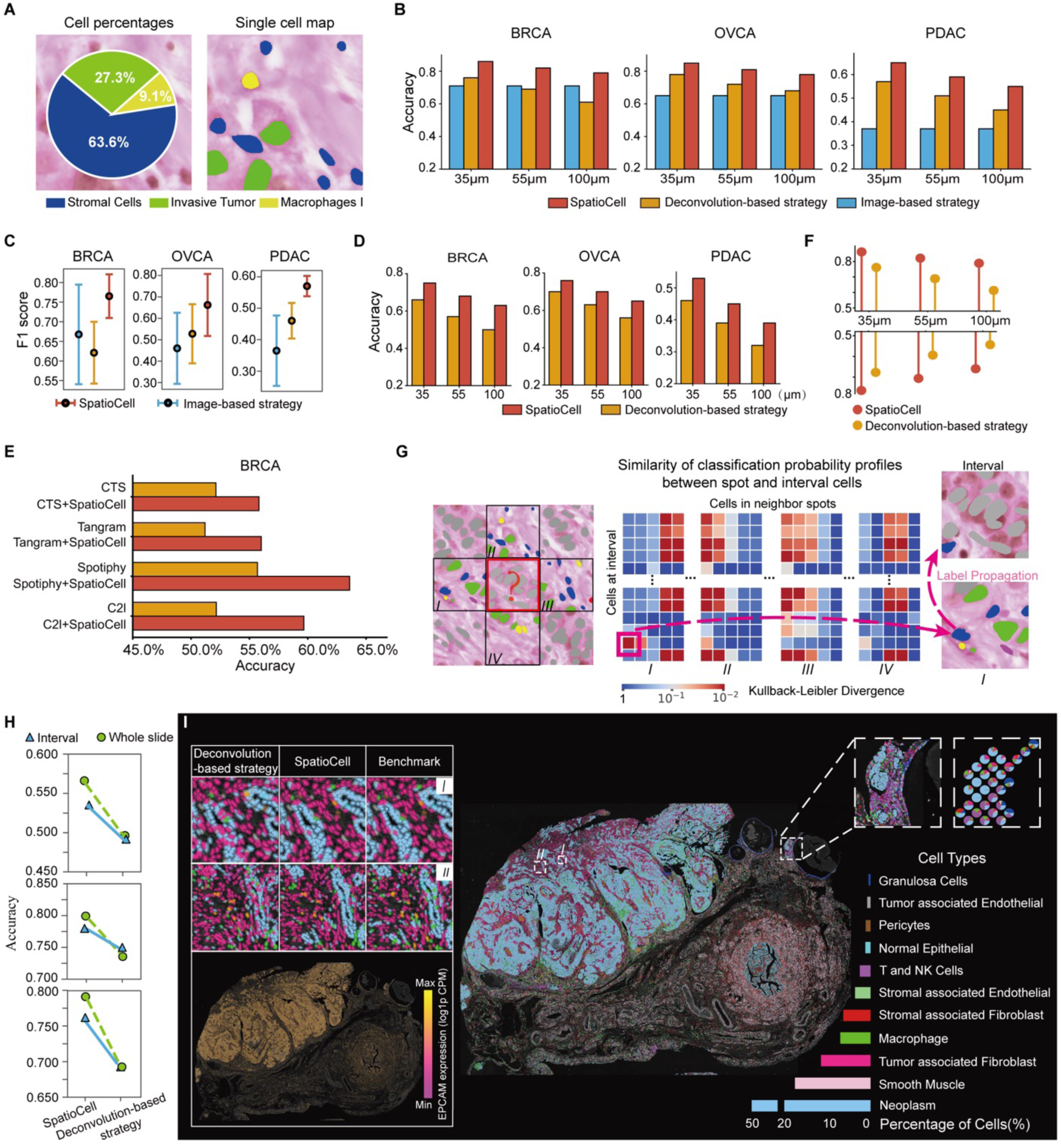
Benchmark and visualization of SpatioCell cell annotation using simulated multicellular-resolution ST data. **(A)**, Schematic of SpatioCell’s cell annotation, which transfers deconvolution-inferred cell type composition into a single-cell spatial map. **(B)**, Comparison of supertype cell annotation accuracy among SpatioCell, deconvolution-based annotation, and H&E classification on simulated datasets (including BRCA, OV, and PDAC tissues) at 35 μm, 55 μm, and 100 μm spot resolutions. **(C)**, Comparison of supertype annotation accuracy (measured by F1-score) among SpatioCell, deconvolution-based methods and H&E classification across three simulated datasets, with circles representing the median and error bars indicating the interquartile range (IQR). Data from the 55 μm spot size is shown. **(D)**, Comparison of subtype annotation accuracy between SpatioCell and deconvolution-based methods on simulated datasets at 35 μm, 55 μm, and 100 μm spot sizes, using ground-truth cell-type compositions as constraints. (**E**), Comparison of within-spot subtype annotation accuracy for deconvolution algorithms alone vs. deconvolution + SpatioCell at 55 μm spot size (CTS, CytoSpace; C2l, Cell2location). **(F)**, Comparison of supertype (top) and subtype (bottom) annotation accuracy within spots across three datasets and various spot sizes between SpatioCell and deconvolution-based methods when interval regions are present. **(G)**, Schematic of SpatioCell’s imputation strategy for spot interval regions. Gray cells represent query cells, and black-framed squares indicate annotated neighboring spots. The heatmap visualizes the KL divergence between query and annotated cells based on probability profiles from module 1, with the darkest red square indicating the lowest divergence. **(H)**, Comparison of supertype annotation accuracy across three datasets at a 55 μm spot size between SpatioCell and deconvolution-based methods, showing accuracy within spot interval regions (blue) and across the entire slide (green). **(I)**, Single-cell spatial map of a simulated OVCA dataset with interval region imputation using SpatioCell. The left panel compares deconvolution-based, SpatioCell, and benchmark annotations (top) and illustrates the inferred EPCAM expression (log1p CPM) distribution generated by SpatioCell (bottom). The top-right inset highlights SpatioCell’s annotation in a lymphocyte-infiltrated tumor region, with deconvolution-derived cell composition pie charts for the same area.

Cell type annotations were tested at different granularities: four broad supertypes and more detailed subtypes. SpatioCell surpassed alternative cell annotation methods that rely solely on a single source of information (**Fig. 3B** and **C**, results of LUCA are in **Fig. S4**). At a commonly adopted 55μm resolution, the annotation accuracy for supertype annotation across four datasets exhibited a significant improvement over the other two strategies. For example, in the BRCA dataset, SpatioCell achieved a supertype annotation accuracy of 82%, improving upon the deconvolution-based and image-based methods by 13% and 10.7%, respectively (**Fig. 3B**). It consistently outperformed both methods across different spot sizes, with advantages more pronounced in larger spot (**Fig. 3B**). For further evaluation on subtype cell identification, we only compared SpatioCell with the deconvolution-based strategy since image-based methods alone lack the resolution needed for precise subtype differentiation. In 55μm-resolution BRCA data, SpatioCell achieved an accuracy of 70.2% across 10 cell types, whereas the deconvolution-based method reached only 59%, demonstrating SpatioCell’s superior ability in fine-grained cell annotation. Similarly, in the OVCA and PDAC datasets, SpatioCell exhibited a comparable boost in cell subtype annotation accuracy (**Fig. 3D**). Furthermore, we integrated SpatioCell with multiple deconvolution algorithms and compared the results to the original methods, showing that SpatioCell consistently improves the accuracy of single-cell assignments **(Fig. 3E)**.

Moreover, we divided simulated spots into capture regions and spot intervals to mimic the lack of sequencing coverage, a common issue in many multicellular-resolution ST. SpatioCell’s performance on capture regions consistently surpassed the deconvolution-based method (**Fig. 3F**). Meanwhile, a key advantage of integrating H&E features is that SpatioCell uses cell-type probability profiles from H&E images (Module 1) to infer cell identities in spot intervals. This is achieved by comparing these profiles with those of neighboring cells within the sequencing spots based on morphological similarity and then transferring cell identity information from these spots to the intervals (**Fig. 3G, Fig. S5A**). To validate SpatioCell’s ability for cell type imputation at interval regions, we measured annotation accuracy across entire tissue sections and within spot interval regions in four datasets. SpatioCell consistently outperformed deconvolution-based methods in both regions (**Fig. 3H, Fig. S5B-C**). Notably, across all four tissues, even with intervals, SpatioCell achieved higher overall annotation accuracy than the deconvolution-based method applied to complete (gap-free) sections (**Fig. S5C**).

As shown in **Fig. 3I**, SpatioCell reconstructs the single-cell distribution in OVCA, effectively revealing its complex TME. SpatioCell more accurately delineates tissue boundaries and tumor structures (such as tubular-shaped tumor) compared to the deconvolution-based method, providing a more structured tissue architecture and cellular spatial distribution. Notably, SpatioCell accurately captured the cell-type distribution at tumor margins (as in **Fig. 3I** *regions I-II*) and identified tumor cells surrounded by T and NK cells adjacent to a follicular cyst, a key feature in the OVCA microenvironment^28^ (**Fig. 3I**). In contrast, deconvolution results provided only a coarse estimate of cell-type proportions, overlooking or blurring critical cellular compositions. Meanwhile, SpatioCell also achieves accurate single-cell–level gene expression mapping based on the high-resolution annotation. As illustrated in **Fig. 3I**, SpatioCell reconstructed a single-cell–resolved expression maps for *EPCAM*, thereby enhancing the spatial resolution of transcriptomic profiles.

### SpatioCell Facilitates Fine-Resolution Mapping of Tissue Architecture Within the Tumor Microenvironment from Multicellular ST

In the TME, biologically significant regions, such as vascular and glandular structures, tumor invasion fronts, and lymphocyte infiltration zones, are often located at the margins where multiple cell types converge. However, these small and spatially intricate structures are difficult to detect or resolve using existing deconvolution methods. To validate SpatioCell’s applicability in revealing these subtle, complex, and crucial structures, we applied it to real multicellular-resolution ST datasets from various tumor samples with spatial spot sizes of 55 µm and 100 µm, representing common sequencing resolutions. In a 100 µm-resolution ST dataset of TNBC, the larger spot size leads to higher cell density per spot (up to 294 cells/spot; **Fig. 4A**), while also capturing more cells for sequencing. The intervals between spots are narrower, with fewer cells in those regions (**Fig. 4B**). In contrast, the 55 µm-resolution 10x Visium ST of OVCA results in fewer cells per spot (**Fig. 4C**). Only 19.5% of the nuclei fall within sequenced regions (**Fig. 4D**). Overall, these datasets, with varying resolutions, cell densities, and interval coverage, provide a comprehensive framework to assess SpatioCell’s adaptability in different scenarios.

**Figure 4.**
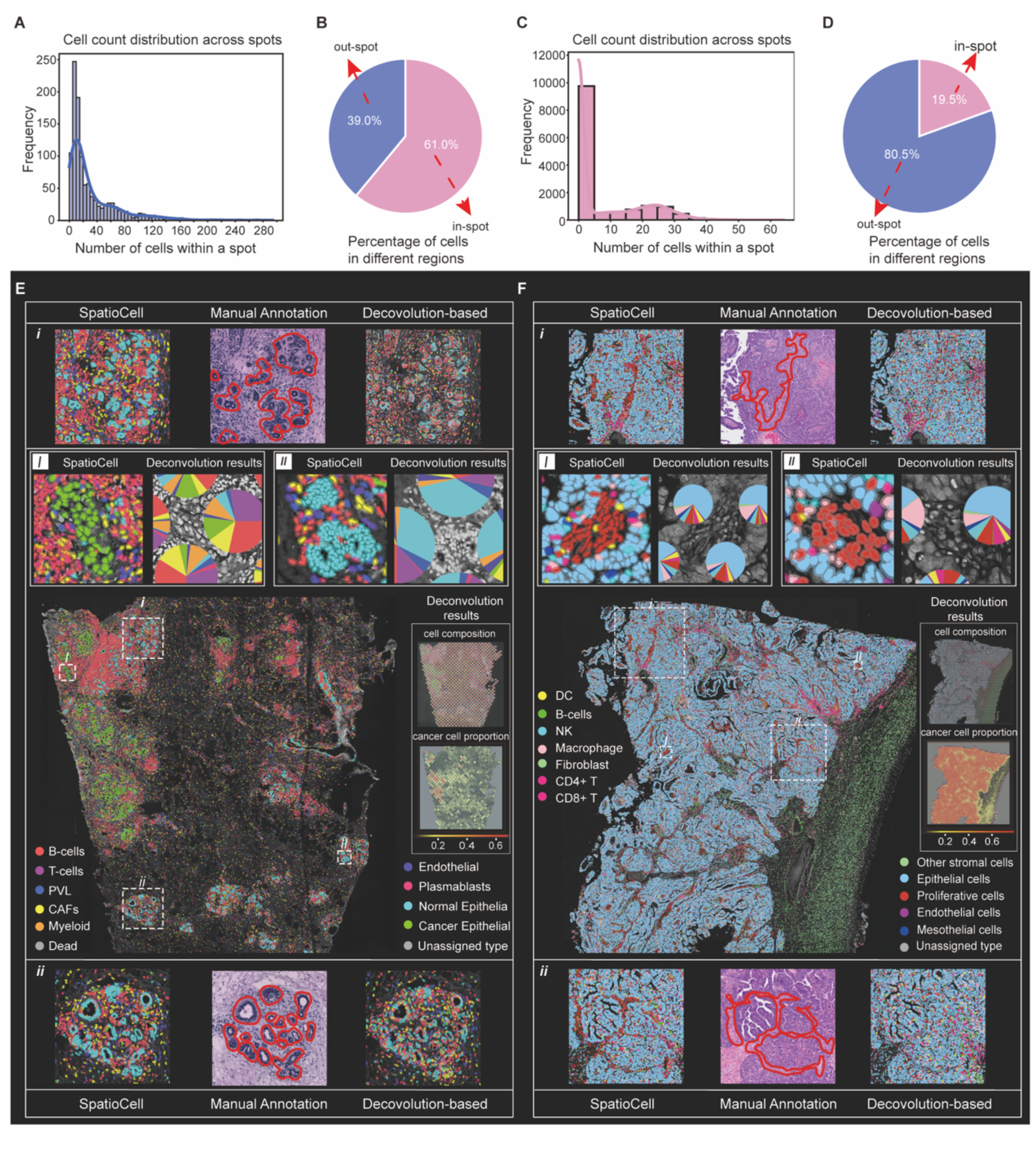
SpatioCell identifies characteristic tissue microenvironment in multicellular-resolution STs. **(A)**, Distribution of cell counts in captured regions of a TNBC sample sequenced with 100 μm spot size, with a density curve showing cell number variability per spot. **(B)**, Proportion of total cell counts between spots and inter-spot regions in the TNBC sample. **(C)**, Distribution of cell counts in captured regions of an ovarian cancer (OVCA) sample sequenced with 55 μm spot size. **(D)**, Proportion of total cell counts between captured and inter-spot regions in the OVCA sample. **(E)**, Single-cell spatial map of the TNBC sample resulted from SpatioCell, with the right panel showing cell composition and cancer cell proportion from deconvolution. (I, II) Comparison of cellular annotation by SpatioCell and deconvolution results, with SpatioCell significantly improving resolution. (i-ii) Comparison of cellular annotation by SpatioCell, manual annotation on H&E, and deconvolution-based methods. **(F)**, Single-cell spatial map of the OVCA sample resulted from SpatioCell, with details similar to (E).

After processing these datasets with SpatioCell, we obtained a comprehensive single-cell spatial atlas of the TNBC data that closely aligned with the manual annotation labeled by a pathologist (**Fig. 4E** and **Fig. S6**). In another TNBC slide, SpatioCell successfully identified a few lymphocyte-enriched regions, with abundant T cells and B cells accumulating at the border between tumor and stroma (**Fig. S7**). In the 55 µm OVCA data, SpatioCell accurately reconstructed conspicuous boundaries between tumor cells and proliferative cells (**Fig. 4F**). However, the deconvolution-based annotation method yielded highly disordered cell-type distributions that concealed these key structural features (**Fig. 4E** and **F**, *i-ii*). These findings confirm that SpatioCell surpasses existing methods in tissue structure reconstruction and spatial organization analysis.

Overall, in these datasets, deconvolution methods only exhibit a rough picture of cell type composition, which limits their ability to perform spatial analysis (**Fig. 4E–F**, panels *I-II*). Prevalent annotation methods, attempt to refine this by integrating segmentation-based cell positions and randomly assigning identities, but they often fail to effectively reconstruct these intricate cellular compositions because of low accuracy. SpatioCell offers superior resolution and accuracy, which establishes the foundation for precise characterization of tumor microenvironment features.

### SpatioCell Enables Deconvolution Errors Correction via a Competitive Balanced Index

Cell-type abundance estimation from deconvolution can be erroneous due to noises introduced by molecular diffusion, sequencing errors, and uneven sampling in ST and single-cell reference data, especially in complex regions. For example, a deconvolution result of a lung squamous cell carcinoma (LUSC) ST data displays widespread tumor cell distribution, even in stromal regions, whereas the H&E image reveals distinct stromal structures and necrotic zones (**Fig. 5A, Fig. S8**). This observation highlights that, by integrating H&E-derived morphological features, SpatioCell not only achieves precise single-cell mapping but also has the potential to refine cell annotations by correcting deconvolution errors. SpatioCell corrects deconvolution-induced errors by calculating Competitive Balance Index (CBI, as detailed in **Methods**), a parameter that quantifies the discrepancy between deconvolution results and H&E-derived nuclear classification. Specifically, when CBI exceeds a preset threshold, SpatioCell progressively refines the deconvolution result using H&E-derived classification probabilities until the discrepancy falls below the threshold.

**Figure 5.**
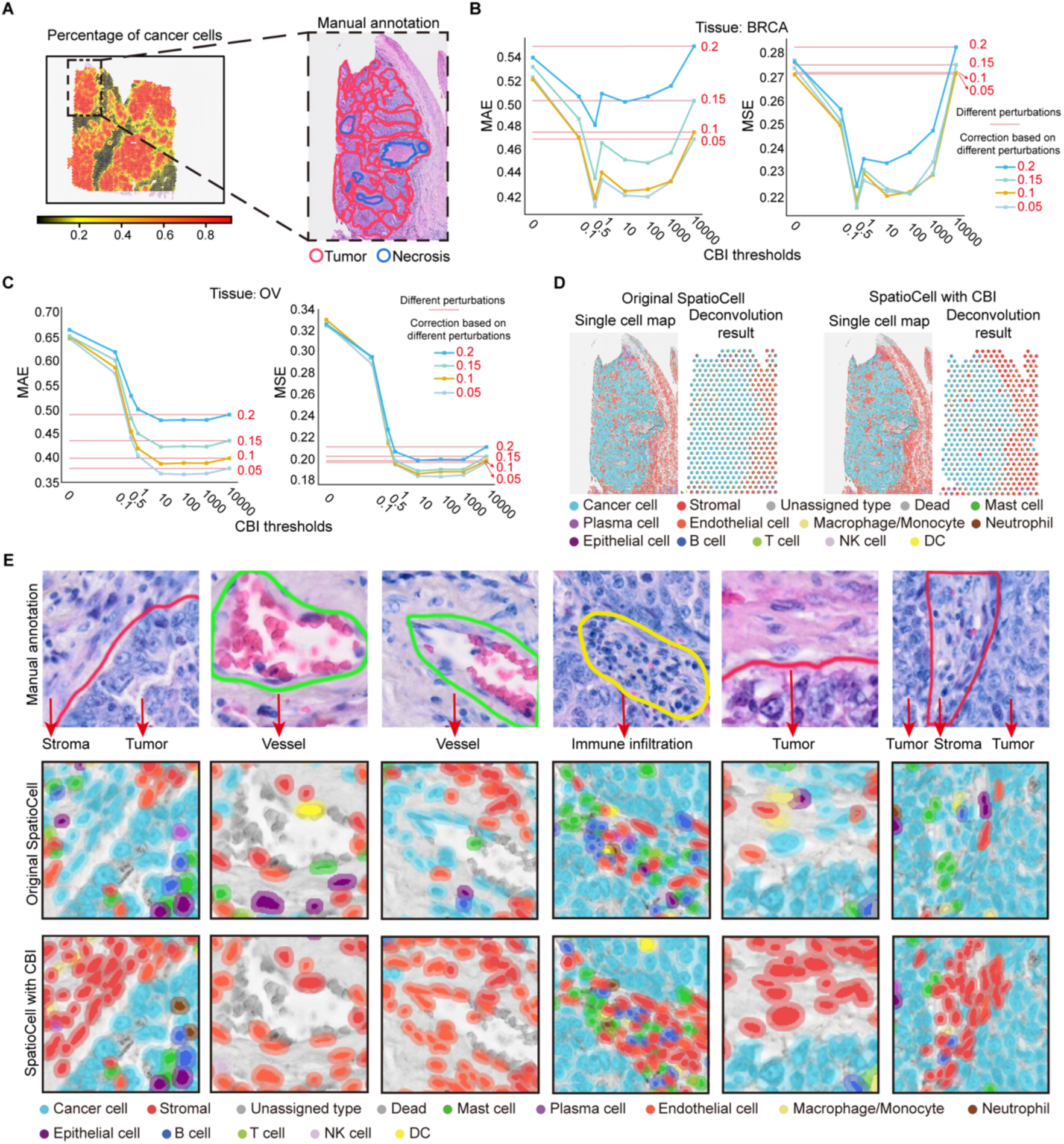
Validation and visualization of SpatioCell with Competitive Balance Index (CBI). **(A)**, Cancer cell proportion estimated by Cell2location in an LUSC sample from 10x Visium (left), with a zoomed-in H&E image (right) highlighting tumor (red) and necrosis (blue) areas as annotated by a pathologist. **(B)**, Distribution of mean absolute error (MAE, left) and mean squared error (MSE, right) of corrected deconvolution results for BRCA ST data across different CBI thresholds. Colored lines represent correction results under various perturbation levels (0.2, 0.15, 0.1, 0.05). Red horizontal lines indicate the corresponding MAE and MSE values without CBI for each perturbation level. **(C)**, Distribution of MAE (left) and MSE (right) of corrected deconvolution results for OVCA ST data across different CBI thresholds, with details similar to (B). **(D)**, Comparison of cell annotation results and deconvolution before and after correction based on CBI. **(E)**, Comparison of cell annotations in 10x Visium-sequenced LUSC ST data: Top row shows manual H&E annotations, second row shows original SpatioCell annotations, and third row shows SpatioCell annotations refined with CBI. The refined annotations align more closely with true tissue structures.

To evaluate the impact of CBI, we introduced controlled errors into the simulated multicellular-resolution ST dataset (described in the previous section) and compared cell composition accuracy before and after CBI-based correction, using mean absolute error (MAE) and mean squared error (MSE). Results show that CBI significantly improves annotation accuracy under various perturbation levels in BRCA and OVCA, with the optimal performance observed at CBI threshold of 0.5 and 10, respectively (**Fig. 5B** to **C**). The improvement observed in OVCA was slightly lower than that in BRCA, likely owing to the lower resolution of its H&E image, half that of BRCA. Our findings suggest that CBI possesses a strong potential for correcting deconvolution errors and that the CBI threshold should be adapted according to the quality of the H&E image. When image quality is high and nuclear classification is reliable, a smaller CBI threshold allows for more drastic correction of deconvolution errors. Conversely, for lower-resolution images, a larger CBI threshold helps preserve annotation stability. In practice, we recommend starting with a higher threshold (e.g., 10).

After applying CBI to the LUSC data above, we observed a notable improvement in the annotation and cell composition of stromal regions embedded within the tumor, which were not clearly delineated in the original deconvolution results (**Fig. 5D**). By correcting deconvolution errors, SpatioCell reconstructs key TME structures, such as small vessels, well-defined tumor borders, and immune infiltrations, closely matching the pathologist’s manual labels (**Fig. 5E**).

Overall, by leveraging CBI, SpatioCell can balance nuclear classification from H&E images with deconvolution-derived composition estimates, allowing for adaptive correction of deconvolution results and revealing key structural features of the TME, thereby enhancing our understanding of TME dynamics. As a result of this correction, single-cell–level cell-type assignments become more accurate, and reliable single-cell–resolved gene expression profiles can be obtained.

### SpatioCell Uncovers Associations between Spatial Heterogeneity and Functional States of CAFs in TNBC

CAFs, as key regulators in the TME, contribute to tumor growth, immune regulation, and metastasis, and show marked heterogeneity in function and spatial distribution^29^. Recent studies have shown that the composition of the cellular neighborhood can influence cell behavior and molecular characteristics through intercellular interactions^30, 31^. However, the associations between CAFs’ spatial neighborhoods and their functional states remain poorly understood, yet deciphering these relationships is crucial for understanding the TME and advancing therapeutic strategies. To address this challenge, SpatioCell provides single-cell–resolved maps of cell types and gene expression, allowing precise analysis of intercellular distances, spatial neighborhoods, and gene expression patterns.

We applied SpatioCell to annotate CAFs in 83 TNBC ST samples^32^ (spot resolution: 100 µm) and stratified them into five spatial neighborhood subtypes based on the compositions of their six nearest neighboring cells (**Fig. 6A**, detailed in **Methods**). Each cluster exhibited distinct neighborhood features, among which Cluster 0 and Cluster 1 showed particularly notable characteristics. Cluster 0 was positioned significantly closer to tumor cells (average ∼22 μm) than other CAF clusters (∼35–40 μm; **Fig. 6B, Fig. S9**), suggesting enhanced capacity to influence local tumor behavior, such as promoting invasion and matrix remodeling^33^. In contrast, Cluster 1 was found nearer to B and T cells (averaging ∼21 μm and ∼41 μm, respectively) than other CAF subtypes (∼43–45 μm to B cells and ∼55–58 μm to T cells; **Fig. 6C**), suggesting a greater likelihood of participating in immune modulation through chemokine signaling. Moreover, Cluster 1 exhibited greater spatial separation from other CAF clusters (**Fig. 6D**), suggesting its localization within a distinct microenvironment. Collectively, these spatial features support functional divergence among CAF subtypes: Cluster 0’s proximity to tumor cells may enable tumor-supportive behavior such as invasion and matrix remodeling, whereas Cluster 1’s closer spatial association with lymphocytes points to a potential role in coordinating localized immune responses.

**Figure 6.**
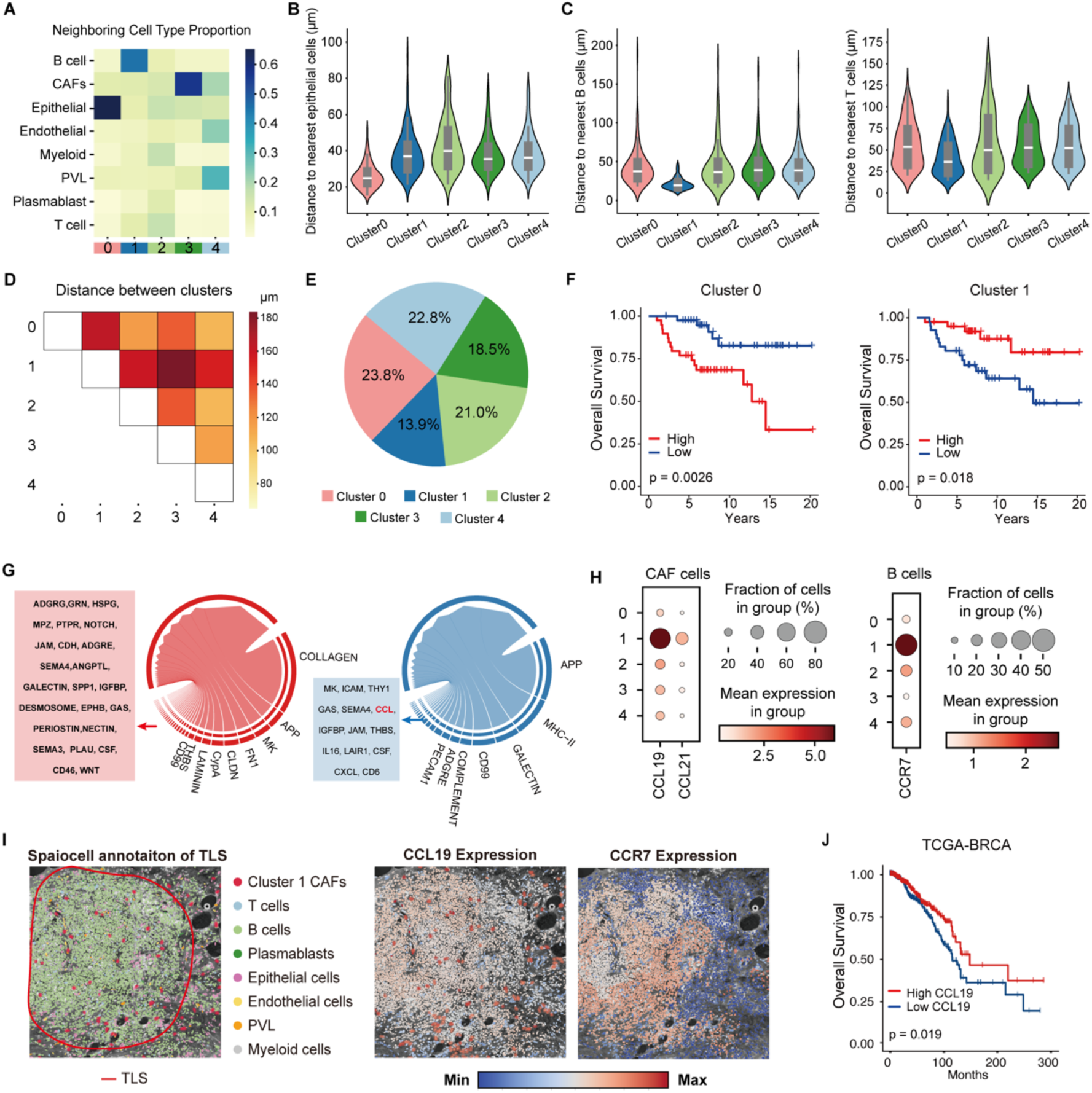
SpatioCell uncovers the spatial heterogeneity and functional states of CAFs in TNBC. **(A)**, Average neighboring cell type composition for each CAF cluster, calculated across all TNBC samples. **(B)**, Comparisons of the average distances from CAFs in each cluster to their six nearest epithelial cells across samples. Distances are measured in microns. **(C)**, Comparisons of the average distances from CAFs in each cluster to their six nearest B cells (left) and T cells (right) across samples respectively. Distances are measured in microns. **(D)**, Heatmap of pairwise spatial distances between CAF clusters. Each value represents the average distance from CAFs in one cluster to their six nearest CAFs in another cluster. **(E)**, Total proportion of CAF clusters across 83 ST samples in the TNBC dataset. **(F)**, Kaplan–Meier survival curves for patients stratified by high (red) vs. low (blue) abundance of CAF Cluster 0 (left) and Cluster 1 (right). High and low groups were defined based on the median proportion of each cluster. P-values were calculated using the log-rank test. **(G)**, Ligand–receptor interaction networks between cells in spots dominated by Cluster 0 (left) and Cluster 1 (right). Band width indicates interaction strength. **(H)**, Inferred cell expression of CCL19–CCR7 axis components across CAF subtypes (CCL19 and CCL21, left) and neighboring B cells (CCR7, right), with dot size indicating the proportion of expressing spots and color intensity representing mean expression per cluster. **(I)**, Single-cell spatial annotation and gene expression mapping of a tertiary lymphoid structure (TLS) region in a TNBC sample resulted from SpatioCell. Left: Spatial distribution of Cluster 1 CAFs and other cell types within and around the TLS. Middle: Spatial distribution of CCL19 expression in the same region. Right: Spatial distribution of CCR7 expression in the same region. **(J)**, Kaplan–Meier survival curve for TCGA-BRCA patients stratified by high (red) vs. low (blue) CCL19 expression. High and low groups were stratified based on the median expression level of CCL19. P-value was calculated using the log-rank test.

Consistent with these spatial-functional distinctions, single-cell atlas statistics showed that the prevalence of each CAF subtype varied across samples, with Cluster 0 being the most abundant and Cluster 1 the least (**Fig. 6E**). Survival analysis further revealed that patients with a higher proportion of CAFs from Cluster 0 had significantly shorter overall survival, whereas those with a higher proportion of Cluster 1 exhibited longer survival durations (**Fig. 6F, Fig. S10-11**). These findings further emphasize the critical role of CAF spatial heterogeneity in modulating CAF function, thereby affecting tumor progression and patient outcomes.

To investigate the intercellular communication between CAF subtypes and their neighboring cells, we classified ST spots according to the dominant CAF subtype within each spot and examined ligand–receptor interactions using CellChat^34^. Cluster 0 showed enrichment of pro-invasive and immunomodulatory interactions, such as FN1-CD44, APP-CD74, PPIA-BSG, and JAG–NOTCH pairs. These interactions drive ECM remodeling, barrier breakdown, immune suppression, and EMT, collectively shaping a tumor-promoting and immune-evasive microenvironment^35, 36, 37, 38^. In contrast, Cluster 1 displayed characteristics of immune-activated CAFs, with strong ligand–receptor signaling activity related to antigen presentation, leukocyte adhesion, and also innate immunity, including HLA-CD4 family, ICAM1/2–ITGAL/ITGB2, and C3–ITGAX/ITGB2 interactions^39, 40, 41^ **(Fig. 6G**).

Building on the precise single-cell annotation, SpatioCell decomposed the spot-level expression and mapped gene expression to each cell location. The gene expression profiles of Cluster 0 and Cluster 1 CAFs were further analyzed at single-cell resolution to explore their distinct roles within the TME. Cluster 0 CAFs showed high expression of immunosuppressive genes, whereas Cluster 1 CAFs were enriched for immune-activating genes (**Fig. S12A**). Noteworthily, Cluster 1 highly expressed CCL19, while its neighboring B and T cells expressed high level of CCR7, a chemokine-receptor pair associated with TLSs and known to guide lymphocyte trafficking via the CCL19/CCL21–CCR7 axis^42^ (**Fig. 6H, Fig. S12B**). This expression profile aligns well with the spatial proximity to B and T cells (∼21 μm and ∼41 μm, respectively) identified by SpatioCell, reinforcing the functional and spatial integration of this immune-modulating CAF subtype. SpatioCell-based annotation and inferred cell expression of CCL19 and CCR7 also revealed that Cluster 1 CAFs are frequently positioned near TLS regions (**Fig. 6I**). Consistent with these findings, analysis of the TCGA BRCA cohort demonstrated that high CCL19 expression was associated with improved survival outcomes (**Fig. 6J**).

In summary, SpatioCell enables single-cell annotation and precise analysis of intercellular distances, spatial neighborhoods, and gene expression patterns, revealing fine-grained spatial heterogeneity of CAFs in TNBC. This analysis uncovers clinically relevant CAF subtypes and underscores the key role of spatial organization in shaping their behavior and influencing tumor progression.

## Discussion

ST offers critical insights into gene expression within the tissue architecture, and the growing accumulation of ST datasets presents valuable opportunities for spatial biology. However, the resolution of most available datasets remains inherently constrained by the size of spatial capture spots. Existing deconvolution methods^17^ ^43^, have sought to infer cell compositions within spots, but these methods lack the spatial precision required to determine the exact localization of individual cells, which makes it difficult to recover detailed structural features and gene expression pattern within the tissue. SpatioCell goes further by integrating imaging data-derived morphological features with transcriptomic measurements to reconstruct single-cell spatial maps from multicellular-resolution ST data. Through detailed analysis of cell distribution, expression and tissue architecture, it holds the potential to advance research in developmental biology, tumor microenvironment studies, and disease pathology.

To achieve this, SpatioCell integrates a novel spatio-morphological learner, re-engineering SAM’s prompting and decoding mechanisms for robust and accurate nuclear segmentation and classification. To evaluate its performance, we conducted tests across six public datasets. SpatioCell outperformed other widely used nuclear segmentation and classification algorithms. Building on this foundation, SpatioCell integrates H&E-derived cell morphology with transcriptomics data, enabling a more biologically meaningful reconstruction of single-cell distributions. Detailly, SpatioCell generates a probabilistic cell-type profile for each nucleus based on its morphology, rather than assigning a hard label. This enables SpatioCell to model the integration as a constrained combinatorial optimization problem, balancing conflicting information from image and sequence data and ensuring a more coherent fusion. In this way, SpatioCell achieves high-resolution single-cell annotation and recovers fine-scale structural details that deconvolution methods often miss. For example, in a 100 µm resolution TNBC ST dataset, SpatioCell vividly restored a normal breast lobule structure heavily infiltrated by T and B cells, alongside neighboring tumor cells. Similarly, in OVCA data, it accurately delineated the tumor-stroma boundary, revealing a stroma-rich region densely populated with proliferative cells, subtle yet biologically significant features otherwise obscured by existing annotation methods.

To validate the usefulness of SpatioCell for ST studies, we use its high-resolution spatial annotation capability to dissect the spatial heterogeneity of CAFs in TNBC. The identification of distinct CAF neighborhood subtypes with divergent transcriptional and functional profiles, along with their characteristic intercellular distances to tumor cells and immune populations, underscores the complex role of spatial context in shaping CAF behavior. Particularly, the contrasting pro-tumorigenic versus immune-activating phenotypes of Cluster 0 and Cluster 1 highlight how CAFs can differently influence tumor progression and immune response depending on their microenvironment. These insights not only advance our understanding of CAF biology but also provide a framework to better integrate spatial distribution with analyses of cellular functional heterogeneity.

SpatioCell marks a significant advancement in ST by integrating image and transcriptomic data to achieve single-cell resolution annotations, enabling a deeper understanding of tissue architecture and its connection to disease. However, its capabilities are still limited by the size of available training datasets for segmentation and classification. Future directions include expanding the training data by annotating more stained images and applying novel computational methods for more precise cell-type classification on H&E images—as demonstrated in ROSIE^44^. Notably, SpatioCell’s success with DAPI-stained images highlights its potential applicability across diverse imaging modalities. With further training, SpatioCell could be extended to subcellular-resolution ST platforms (such as Stereo-seq^10^ and Xenium^45^), where nuclear contours (e.g., DAPI-stained) are readily available, enabling more precise annotations in subcellular and single-molecule resolution contexts. These advancements expand SpatioCell’s applicability to all spatial omics platforms and further drive the translation of spatial omics into biomedical research.

## Materials and Methods

### Module I of SpatioCell: H&E image segmentation and analysis

The H&E processing module is designed to achieve precise nuclear segmentation and analysis on H&E-stained images. The algorithm is implemented in Python, based on the PyTorch deep learning library and the MMSegmentation framework. It outputs nuclear segmentation masks, centroid and contour coordinates, and cell type classification probability profiles. Pretrained weights for our segmentation and classification models are available for download, allowing users to perform direct inference on new H&E images.

#### Architecture of Prompter

The encoder adopts the MixTransformer-b3 (mit-b3) backbone from the Segformer, consisting of an Overlap Patch Embedding layer followed by four hierarchical Transformer blocks. Each Transformer block includes Efficient Self-Attention, a Mix-Feedforward Network (Mix-FFN), and Overlap Patch Merging, which together extract multi-scale features while preserving local continuity and spatial resolution. Feature maps at each level are downsampled progressively (1/4, 1/8, 1/16 and 1/32). The extracted feature maps are first processed through 1×1 convolutions and upsampled to the highest resolution. They are then concatenated along the channel dimension and passed into two decoder branches: the prompt branch generates heatmaps of nuclear centroids, while the classification branch produces semantic segmentation maps representing cell-type probabilities. Both branches employ multiple convolutional layers to capture multi-scale contextual information. The resulting features are concatenated and fed into their respective decoder heads.

In the prompt head, high-value points in the heatmap are extracted as prompt points for SAM. In the classification branch, a softmax layer is applied to generate class probability profiles, with each value indicating a cell-type probability. The Prompter thus provides both spatial prompts for SAM and type-specific probabilities for downstream annotation.

#### Architecture of SAM

SpatioCell employs SAM for nuclear segmentation. SAM includes an image encoder (ViT), a prompt encoder, and a mask decoder. The image encoder extracts global features from image patches with positional encoding and self-attention. Prompt embeddings, generated from nuclear centroids by the prompter, are fused with image features in the decoder through self- and cross-attention mechanisms. The decoder refines these features using MLPs and produces segmentation masks by integrating them with upsampled image features. Full architectural details are provided in **SI Note 2**.

#### Prompter Training

SpatioCell adopts a two-stage training strategy, in which the prompter and SAM are trained independently. The Prompter is optimized through two supervised learning objectives: nuclear centroid localization and cell-type classification. To enable accurate localization of nuclear centroids, the prompt branch is trained using Gaussian kernel-based heatmaps as supervision, which are generated on the heatmap to represent the probability of a pixel being at the center of a nucleus. Owing to the imbalanced distribution on the heatmap, we used Focal loss in this branch, defined as:

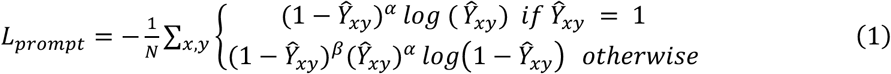

where Ŷ_xy_ is defined as the heatmap predict value of pixel (x, y), α and β are hyperparameters, and N denotes the number of total prompt points.

The classification branch is trained as a semantic segmentation task, and the total loss is defined as the sum of Dice loss and cross-entropy loss:

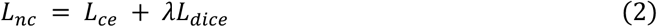

where λ represents the hyperparameter that weighs the impact of the two losses, and in this study, λ = 3. Here, we use the mit_b3 weight pretrained on the ImageNet dataset^19^. Additional details are available in **SI Note 2**.

#### SAM fine-tuning

To improve its performance on nuclear segmentation, SAM was fine-tuned on a nucleus instance segmentation dataset. For each nucleus, the coordinates of one pixel within the ground truth mask were randomly selected as a positive prompt. Additionally, with a probability of 50%, a negative prompt was added by selecting the coordinates of a pixel from the nearest neighboring nucleus. This strategy helps the model better distinguish between closely adjacent nuclei. The predicted mask was trained using a combination of Dice loss and Focal loss between the predicted and the ground truth mask:

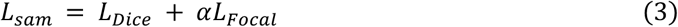

where α=1 is set as the default value during training.

#### Nuclear segmentation inference

During inference, the Prompter generates a predictive heatmap of nuclear centroids along with a semantic map indicating nuclear types. Pixels with heatmap values exceeding a threshold τ are selected as positive prompts, and their corresponding locations in the semantic mask are used to assign cell type labels. The selected prompts, along with the input image patches, are passed to SAM for instance segmentation. Finally, overlapping predictions resulting from the sliding-window tiling strategy are filtered using Matrix NMS. Additional details are available in **SI Note 2: *Inference and Postprocessing***. **Module II of SpatioCell: Single-cell resolution annotation and gene expression mapping of ST *Estimating cell type compositions within spots***

The cell annotation module is compatible with outputs from various spatial deconvolution tools (e.g., Cell2location, Spotiphy, Tangram). In this study, Cell2location was employed using scRNA-seq data derived from the same tissue type. Counts from ST data are preprocessed with mitochondrial and uninformative genes were filtered out. Median number of segmented nuclei within spots was used in ‘N_cells_per_location’, and ‘detection_alpha’ was set to 200. All other parameters were kept at their default settings. Deconvolution results are provided in **Fig. S6** to **8**. Details on implementation of other deconvolution algorithms are available in **SI Note 3: *Deconvolution algorithms applied in benchmarks***. ***Cell type annotation within spots***

Cell type annotation within a spot involves two primary inputs: (1) cell nuclei center coordinates and cell type probability distribution, which are output by SpatioCell’s H&E image processing module, and (2) cell type compositions within the spot, which are provided by deconvolution approaches. These two inputs are then combined to estimate the distribution of cell types across different spots in a tissue section.

Step 1: Estimating the number of cells in each spot

Each spot is defined by its spatial location and size, which enables the calculation of the total number of cells it contains. Let the total number of cells within a given spot be *N_spot_*, and the corresponding cell type proportions obtained from deconvolution be represented as *P_i_* for each cell type *i*. The total number of cells of each type in the spot can be estimated as:

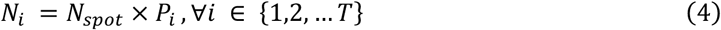

where *n_i_* represents the estimated number of cells of type *i* in the spot. Step 2: DP for cell type assignment After estimating the number of cells of each type within a spot, we aim to assign these cell types to each cell such that the total sum of assigned probabilities across all cells within the spot is maximized. This problem can be formulated as a constrained combinatorial optimization task (a variant of the knapsack problem), where each cell in a spot is considered an “item”, associated with cell type probability profile as “values”.

SpatioCell performs the cell annotation through a DP algorithm that recursively searches all feasible label assignments (i.e. states) across nuclei to identify the globally optimal outcome. Let *P*_*n,i*_ denote the probability that nucleus n belongs to cell type *i*, the DP algorithm proceeds as follows:

(1) State Space Initialization: Define the state space for the cell type assignment problem. The state *s* is represented as a vector of cell counts for each type, i.e. *s* = (*n*_1_, *n*_2_, . . ., *n*_*T*_), *n*_*i*_ ≤ *N*_*i*_ where *n*_*i*_ is the number of cells of type *i* that have been assigned so far.
(2) Transition Between States: For each state *s*, the algorithm explores all possible directions of state transitions, *e*^(*i*)^. For each transition, the goal is to maximize the sum of the probabilities of assigning a cell to a particular type while respecting the total number of cells for each type, as constrained by the deconvolution results. The transition function is given by:

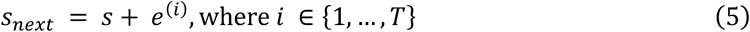

and for each transition, the objective is to maximize the total probability:

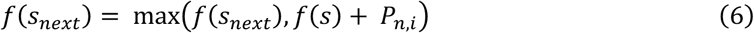

where *f*;(*s_next_*) represents the current maximum score for a given state and *P_n,i_* is the probability of the n^th^ cell belonging to type *i*.
(3) Backtracking the Optimal Path: The DP starts from *s*_0_ = (0, 0, . . .,0), and iteratively updates states until reaching s_final_ = (*N*_1_, *N*_2_, . . ., *N*_*T*_). Once all possible transitions have been explored, the optimal assignment is found by backtracking from the final state to the initial state, determining the optimal direction (i.e., cell type assignment) at each step.

#### Cell type imputation in spot interval

Segmented cells are first located within each interval region based on spot coordinates and size. For each interval, neighboring spots are identified, and cell types are assigned by comparing type probability profiles between interval and neighboring cells. The most frequent type among the top-k most similar neighbor nuclei is selected (k is set to 5 in the study). Similarity is measured using KL divergence as follows:

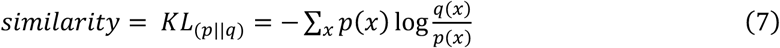

where *p*(*x*) and *q*(*x*) denote cell type probability profile of the query cell at interval and annotated cell in neighbor spots respectively.

#### Gene expression mapping at single-cell level

Gene expression within individual spots was decomposed into cell-type–specific contributions following the strategy implemented in Spotiphy^17^. It assumed that for each spot *s* and gene *g*, the observed counts *x*_*s,g*_ was modeled as a multinomial distribution across cell types *t*:

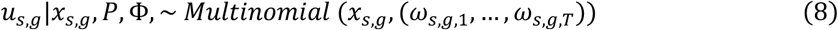

where the allocation weight across cell types *t* is defined by:

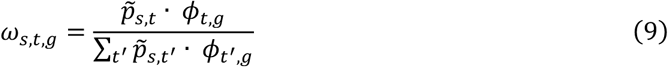

Here, *p̃_s,t_* denotes the proportion of cell type *t* in spot *s* estimated by SpatioCell, and ϕ_*t,g*_ represents the expression profile of gene *g* in cell type *t* obtained from scRNA-seq data. The expected expression assigned to each cell type is then given by:

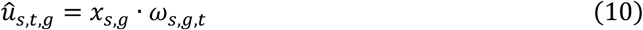

The inferred cell-type–specific expression profiles were precisely assigned to individual cells according to the single-cell annotations derived from SpatioCell, thereby generating a high-resolution spatial map of cell identities and gene expression. For each cell at interval, gene expression was imputed by borrowing information from the spatially nearest neighbor cell whose morphological probabilistic profile exhibited the highest similarity.

### Competitive Balance Index (CBI) for deconvolution improvement

CBI quantifies the relative discrepancy between image-based nuclear classification and deconvolution-derived cell-type assignments within a spot. Specifically, after annotation, we compute the total cell-type probability (*P_assigned_*) and compare it to the sum of the maximum class probabilities of nucleus (*P_images_*), defined as:

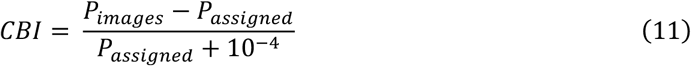

where 10^−H^ is added as smoothing factor to prevent exaggerated CBI values.

If the discrepancy exceeds the predefined CBI threshold, SpatioCell iteratively updates cell-type assignments by identifying nuclei whose H&E-based type differ from the deconvolution-based assignment. These cells are ranked by their highest classification probability across all types, and their assigned supertypes are replaced with H&E-predicted types progressively. After each update, *P_assigned_* is recalculated, and the process continues until the CBI falls below the threshold.

Subtypes for adjusted cells are further assigned from the most similar unmodified cell within the same spot that shares the same supertype (similarity defined as in interval imputation).

### Benchmarks on nuclear segmentation and classification

We benchmarked the image processing module of SpatioCell with five algorithms including Hover-Net, Cellpose3, StarDist, Mask R-CNN and CellProfiler, on six public datasets: Kumar, CPM 17, PanNuke, CoNSeP, MoNuSAC, and CellBinDB. Nuclear segmentation performance was evaluated on all except MoNuSAC dataset since StarDist was pretrained on it. Cell type classification was benchmarked on PanNuke, CoNSeP, and MoNuSAC to compare SpatioCell, Hover-Net and Mask R-CNN.

For each dataset, we used its original training, validation, and testing splits. In PanNuke dataset, we conducted a three-fold cross-validation on its predefined partitions to ensure robust evaluation. We used AJI, DQ, SQ and PQ to comprehensively validate nuclear instance segmentation. AJI and SQ measure the segmentation quality while DQ evaluates models’ ability in nuclei detection (F1-score for detection). PQ considered SQ and PQ together. For cell type classification, we used mPQ and F_C_, defined as the product of the detection F1-score and overall classification accuracy for correctly detected nuclei.

Segmentation performance on DAPI-stained images is tested using 10x Genomics dataset in CellBinDB. Since SpatioCell, Hover-Net, and Mask R-CNN do not offer pre-trained weights for DAPI-stained images, we trained these models from scratch using DAPI-stained images from stereo-seq dataset in CellBinDB. In contrast, Cellpose3 and StarDist were evaluated using their respective pre-trained weights, “cyto3” and “fluoro”. Additional details in the ***compared methods***, ***image preprocessing***, ***dataset*** and ***evaluation metrics* are available in SI Note 1**.

### Evaluation of annotation using Xenium data

To simulate multicellular-resolution ST data, we obtained Xenium datasets from 10x Genomics website, including BRCA, OVCA, PDAC, and LUCA samples. Each dataset provides H&E and DAPI images, DAPI-based nuclear information (regarded as ground truth nuclei), affine transformation matrices and cell annotations (except for LUCA, which we generated marker-based annotations using Seurat). H&E images were aligned to DAPI by ‘*skimage*’ according to affine transformation matrices and then partitioned into 35, 55, and 100 μm square to simulate spots. Squares located at odd rows with even columns and even rows with odd columns were designated as intervals. Segmented and ground truth nuclei were assigned to a spot based on their centroid coordinates. Simulated deconvolution results were generated by calculating the proportion of annotated cell types within each spot based on the provided annotations.

To evaluate annotation accuracy, we used a mutual nearest neighbor (MNN) strategy to align segmented and ground truth nuclei based on their centroids. Cell pairs identified as mutual nearest neighbors were considered aligned. Unmatched ground truth nuclei and uncertain labels were excluded from computing accuracy. Additional details in ***Xenium Human Lung Cancer Cell Type Annotation*** and ***Mapping H&E image segmented cells with Xenium annotation*** are available in **SI Note 3**.

### Evaluation of CBI impact on deconvolution results using Xenium data

We used BRCA and OVCA datasets to evaluate robustness by introducing perturbations into the simulated deconvolution results within each spot. Specifically, 5%, 10%, 15%, 20%, and 30% of cell assignments were randomly perturbed per spot (rounded down, with at least one cell modified per spot). For each perturbation level, we applied CBI-based correction using thresholds of 0, 0.1, 0.5, 1, 10, 100, 1000, and 10000 (representing no correction), and compared the corrected compositions and the perturbed compositions against the ground truth. MAE and MSE were used as evaluation metrics, defined as follows:

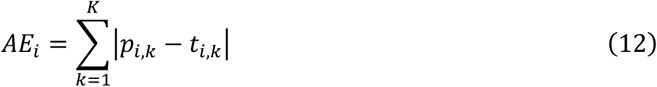

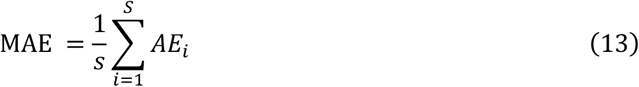

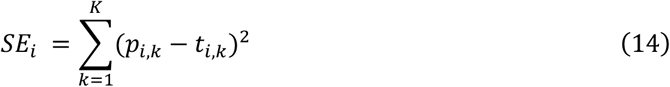

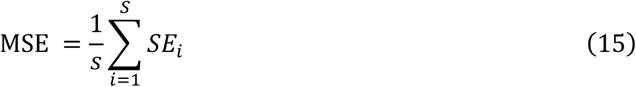

where *p*_*i,k*_ and *t*_*i,k*_ denote the estimated and ground truth proportion of cell type *k* in spot *i* respectively.

### CAF neighborhood clustering and spot classification

We analyzed ST data from 83 breast cancer patients with ER-, PR-, and HER2-status from the Wang et al. cohort^32^. Single-cell annotation of the ST data was performed using SpatioCell. For each annotated cell, the six nearest neighboring cells within a 20 μm radius were identified, and the cell-type compositions were recorded to construct a local neighborhood profile.

Focusing on CAFs, we applied K-means clustering to six-cell neighborhood profiles and identified five spatial subtypes based on the elbow plot.

For downstream transcriptomic analysis, we quantified the number and proportion of each CAF neighborhood subtype within every ST spot. Each ST spot was assigned a dominant CAF subtype if over 80% of its CAFs belonged to the same cluster. Only these subtype-dominant spots were retained, while spots without a clear dominant population were excluded to avoid confounding effects from cellular heterogeneity.

### Spatial proximity measurements

To characterize the spatial proximity between CAF clusters and key cellular components of the tumor microenvironment, two types of distances were calculated based on cell centroids using scikit-learn’s *‘NearestNeighbors’*: (1) CAF-to-cell-type distances—for each CAF in a given cluster, we computed the average distance to its six nearest neighboring B cells, T cells, or epithelial cells, respectively; and (2) inter-cluster distances—to assess spatial relationships between CAF clusters, we calculated the average minimum distance from cells in cluster i to their six nearest neighbors in cluster j. The final pairwise distance matrix was obtained by averaging across samples.

### Inferring cell communication networks and analyzing cell neighbors

Cell communication analysis was conducted using the R package *‘CellChat’* (version 2.2.0) on ST data stratified by dominant CAF subtypes. Intra-cluster ligand–receptor interactions were visualized using the ‘*netVisual_chord_gene*’ function, and only interactions with a communication probability greater than 0.01 were included in the chord diagrams.

### Survival analysis

The *‘survival’* (version 3.8.3) and *‘survminer’* (version 0.5.0) R packages were used to perform survival analysis. For each CAF neighborhood subtype and for CCL19 expression (in TCGA-BRCA), patients were divided into high and low groups based on median values. Kaplan–Meier survival curves were generated using the *‘survfit’* function, and differences in overall survival between groups were evaluated using the log-rank test via the *‘survdiff’* function. Visualization of survival curves was performed using the *‘ggsurvplot’* function.

### Statistical analysis

The highly differential genes in each cluster of spatial transcriptomics data were identified using the Python package *’scanpy’* and tested using Wilcoxon rank-sum test. Differences in overall survival in both TNBC cohort and TCGA cohort were evaluated using the log-rank test. All tests were two sided, and p values < 0.05 were considered statistically significant.

### Paper Writing

To improve the clarity and fluency of the manuscript, we utilized OpenAI’s ChatGPT-4o model for English language polishing. The model assisted in refining grammar, style, and readability, while all scientific content was reviewed and validated by the authors.

## Data availability

All data used in this study were previously published and publicly available. CPM17 and Kumar datasets are available from https://github.com/vqdang/hover_net, CoNSeP dataset and PanNuke dataset can be downloaded from https://warwick.ac.uk/fac/cross_fac/tia/data/. MoNuSAC dataset is available from https://monusac-2020.grand-challenge.org, CellBinDB dataset is available from https://db.cngb.org/search/project/CNP0006370.

Xenium datasets of FFPE human breast cancer, FFPE human lung cancer, FFPE human ovarian cancer, and FFPE human pancreatic cancer are downloaded from 10x Genomics (https://www.10xgenomics.com/products/xenium-in-situ/preview-dataset-human-breast; https://www.10xgenomics.com/datasets/ffpe-human-lung-cancer-data-with-human-immuno-oncology-profiling-panel-and-custom-add-on-1-standard; https://www.10xgenomics.com/datasets/xenium-prime-ffpe-human-ovarian-cancer; https://www.10xgenomics.com/datasets/pancreatic-cancer-with-xenium-human-multi-tissue-and-cancer-panel-1-standard).

Visium datasets of human ovarian cancer and lung cancer are available on the 10x Genomics websites (https://www.10xgenomics.com/datasets/human-ovarian-cancer-11-mm-capture-area-ffpe-2-standard; https://www.10xgenomics.com/datasets/human-lung-cancer-ffpe-2-standard). The TNBC ST data and the associated brightfield H&E images can be downloaded via Zenodo: https://doi.org/10.5281/zenodo.8135721.

The human breast cancer scRNA-seq dataset is available from the Gene Expression Omnibus under accession number GSE176078 (https://www.ncbi.nlm.nih.gov/geo/query/acc.cgi?acc=GSE176078).

The human ovarian cancer scRNA-seq dataset is available on Mendeley Data (https://doi.org/10.17632/rc47y6m9mp.1). The non-small cell lung cancer scRNA-seq dataset is available on Zenodo (https://zenodo.org/records/7227571).

## Acknowledgments

We are grateful to Dr. Zaoyu Wang and Dr. Dengfeng Cao from the Department of Pathology, Renji Hospital, for their expert guidance and contributions to the annotation of histopathological slides.

## Author Contributions

Conceptualization: J.S. Methodology: N.H., ZR.W., and J.S. Investigation: N.H., ZR.W., Y.Y., Z.Z., F.X., and ZY.W. Visualization: N.H., Y.Y., Y.Z., and J.W.

