## Supplementary material for "SpatioCell: Deep Integration of Histology and Spatial Transcriptomics for Profiling the Cellular Microenvironment at Single-Cell Level": SI (from another version of this preprint: version 1, 2025-04-12)

**Supplementary Information for**

### SpatioCell: A Deep Learning Algorithm for High-Resolution Single-Cell Mapping through Deep Integration of Histology Image and Sequencing Data

Naiqiao Hou[#1,2], Zhaorun Wu[#5], Yue Yu[#1,2], Yanqi Zhang[1,6,7], Jiayun Wu[1,2], Zhixing Zhong[5], Fangfang Xu[5], Zeyu Wang[2], Chaoyong Yang*[2,5], Weihong Tan*[2,3,4], Jia Song*[1,2]

[1]Institute for Developmental and Regenerative Cardiovascular Medicine, MOE-Shanghai Key Laboratory of Children's Environmental Health, Xinhua Hospital, Shanghai Jiao Tong University School of Medicine, Shanghai 200092, China

[2]Institute of Molecular Medicine, Renji Hospital, School of Medicine, Shanghai Jiao Tong University, Shanghai 200127, China.

[3]Molecular Science and Biomedicine Laboratory (MBL), State Key Laboratory of Chemo/Biosensing and Chemometrics, College of Chemistry and Chemical Engineering, College of Biology, Aptamer Engineering Center of Hunan Province, Hunan University, Changsha, Hunan 410082, P. R. China.

[4]Zhejiang Cancer Hospital, Key Laboratory of Zhejiang Province for Aptamers and Theranostics, Hangzhou Institute of Medicine (HIM), Chinese Academy of Sciences; Hangzhou, Zhejiang 310022, P. R. China.

[5]Institute of Artificial Intelligence, Department of Chemical Biology, College of Chemistry and Chemical Engineering, Xiamen University, Xiamen, 361102, China.

[6]School of Life Sciences, Inner Mongolia University, Hohhot, 010000, China

[7]School of Life Sciences, Fudan University, Shanghai, 200438, China

These authors contributed equally: Naiqiao Hou, Zhaorun Wu, Yue Yu

**8 Supplementary Figures**

### Supplementary Figures

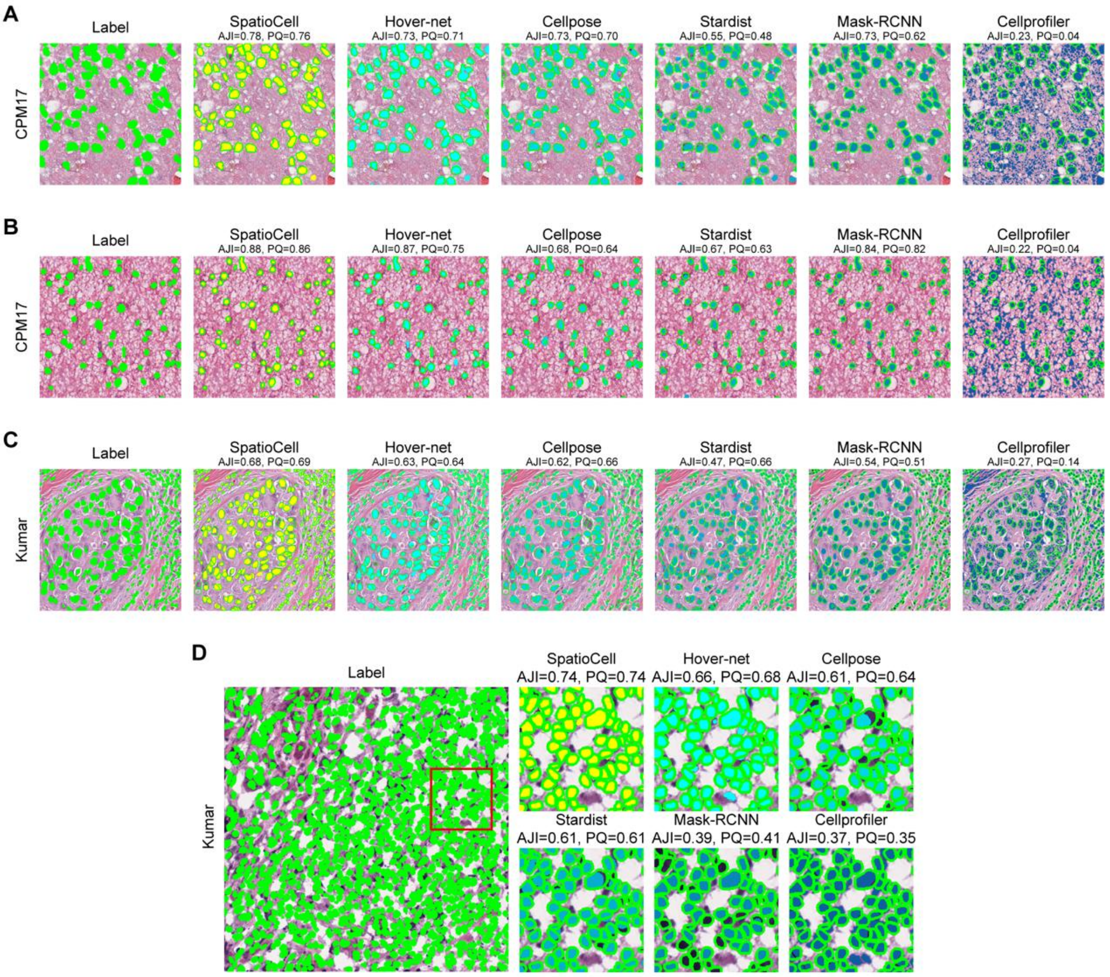


**Supplementary Figure 1. Visualization of SpatioCell's segmentation on Kumar and CPM17.** A-D. Visualized results of nuclei segmentation performance of SpatioCell, Hovernet, Cellpose, Stardist, Mask-RCNN and Cellprofiler on CPM17 and Kumar.

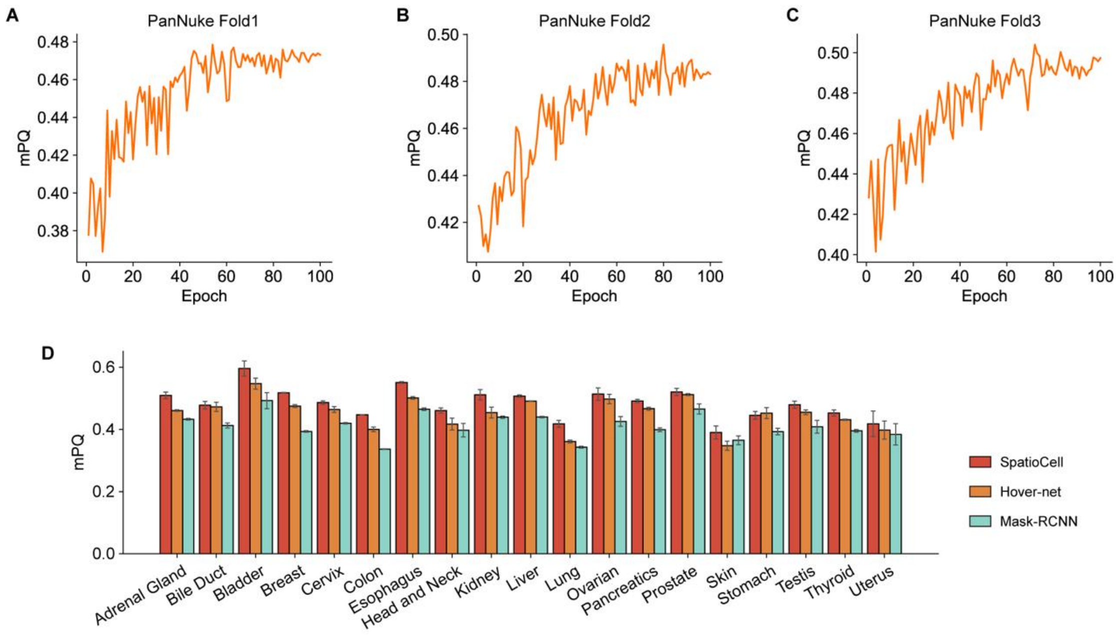


**Supplementary Figure 2. Average segmentation performance of SpatioCell across cell types on PanNuke.** A-C. mPQ score of validation set in PanNuke dataset during training process. D. mPQ of SpatioCell on PanNuke dataset on 19 types of tissues, achieving the best performance in 18 of 19.

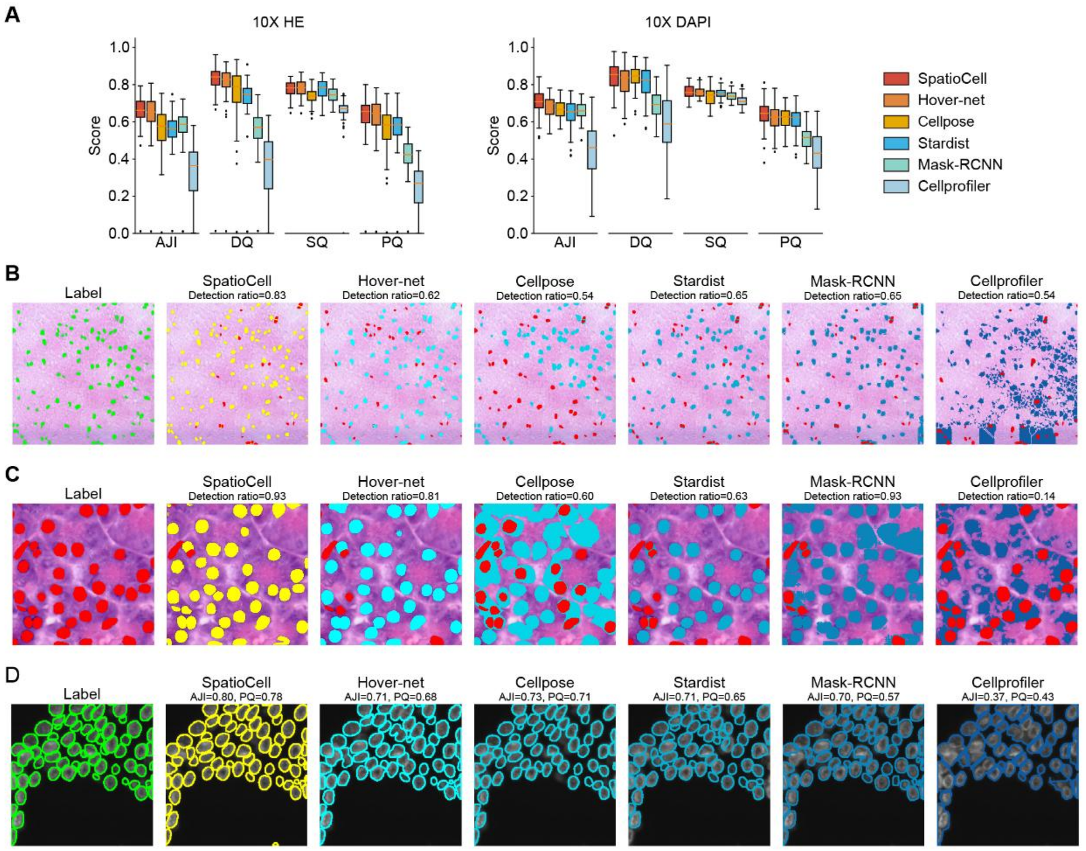


**Supplementary figure 3. Nuclear segmentation performance of SpatioCell on CellbinDB.** A. Nuclear segmentation performance of different methods on H&E-stained images of FFPE samples and DAPI-stained images from 10x Genomics in the CellbinDB dataset. B, C. Visual samples of nuclear segmentation on H&E stained FF and FFPE samples from CellbinDB, with detection ratio annotated. D. Visual samples of nuclear segmentation on a 10X Genomics DAPI image in CellbinDB, labeled with AJI and PQ scores.

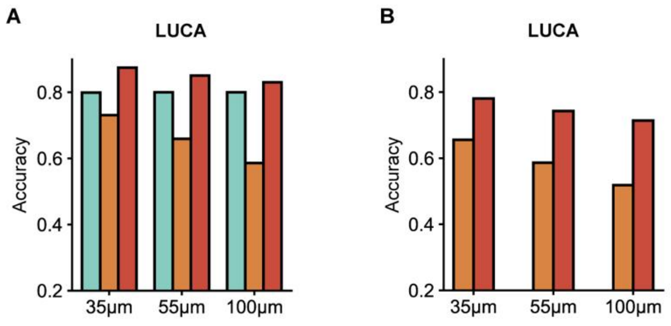


**Supplementary figure 4. Annotation accuracy of SpatioCell using simulated lung cancer.** A. Comparison of supertype cell annotation accuracy between SpatioCell, deconvolution-based annotation, and H&E classification on simulated LUCA dataset at 35 μm, 55 μm, and 100 μm spot resolutions. B. Comparison of subtype annotation accuracy between SpatioCell and deconvolution-based methods on simulated LUCA dataset at 35 μm, 55 μm, and 100 μm spot sizes.

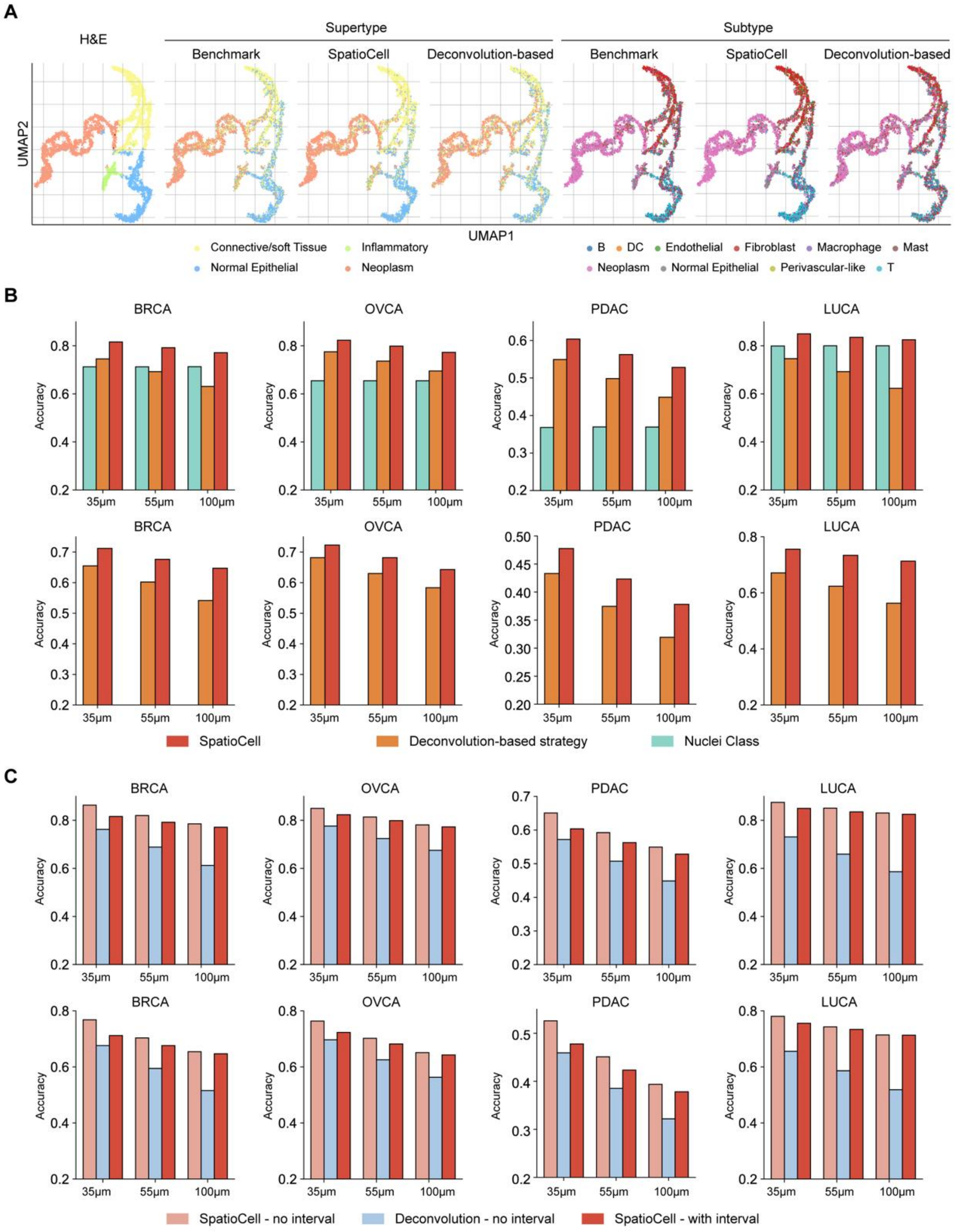


**Supplementary figure 5. Interval imputation performance of SpatioCell in Xenium-simulated multicellular resolution data**. A. Umap of probability profiles generated by the nuclei classification model, colored by cell types in nuclei classification, minor types and major types. B. Wholeslide annotation accuracy of SpatioCell for supertype (top) and subtype (bottom) with intervals on simulated datasets, including BRCA, OVCA, PDAC and LUCA. C. Supertype (top) and subtype (down) Cell annotation accuracy of SpatioCell in simulated datasets with and without intervals, compared with deconvolution-based method in datasets without intervals.

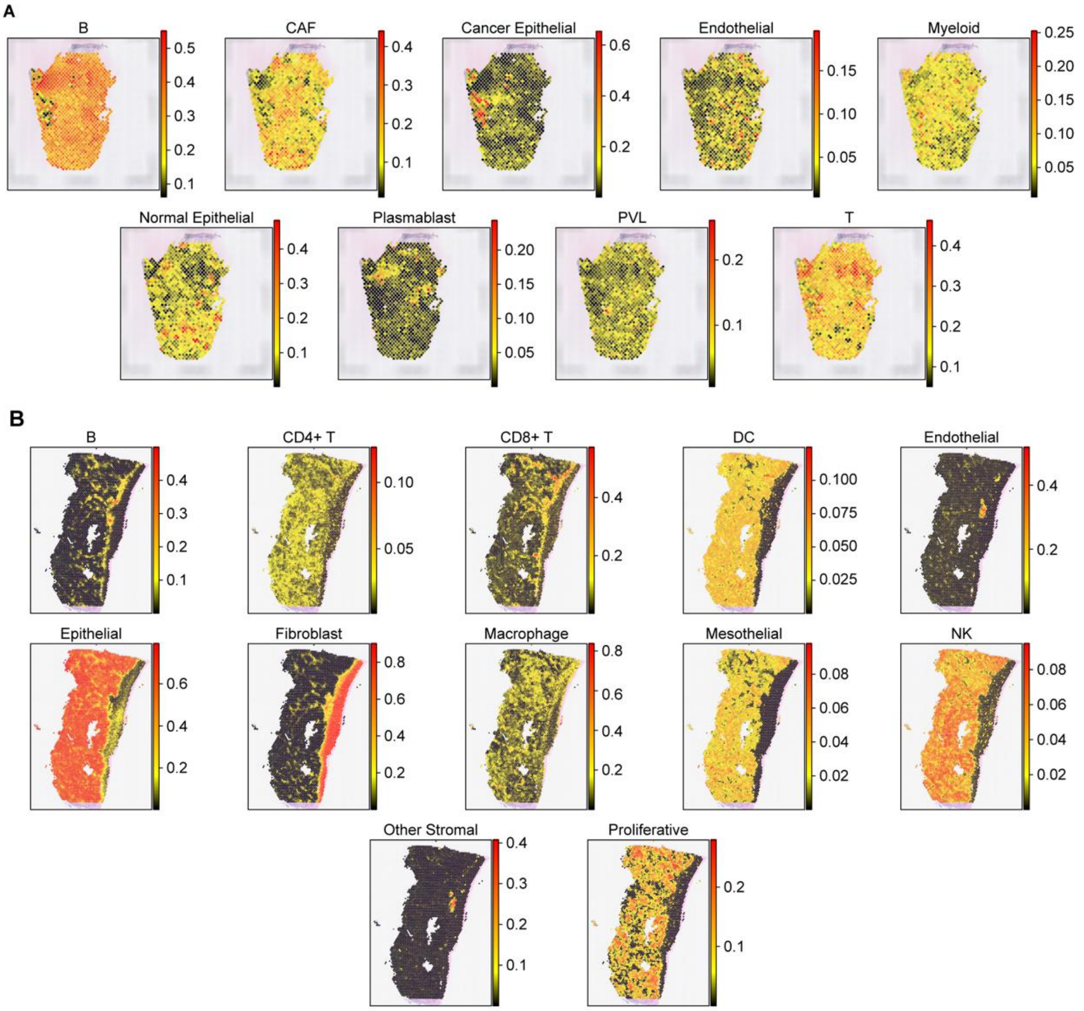


**Supplementary Figure 6. Deconvolution results of TNBC P10 and OVCA.** A, B. Cell abundance of each cell type in TNBC P10 and OVCA spatial transcriptomics data deconvoluted by Cell2location.

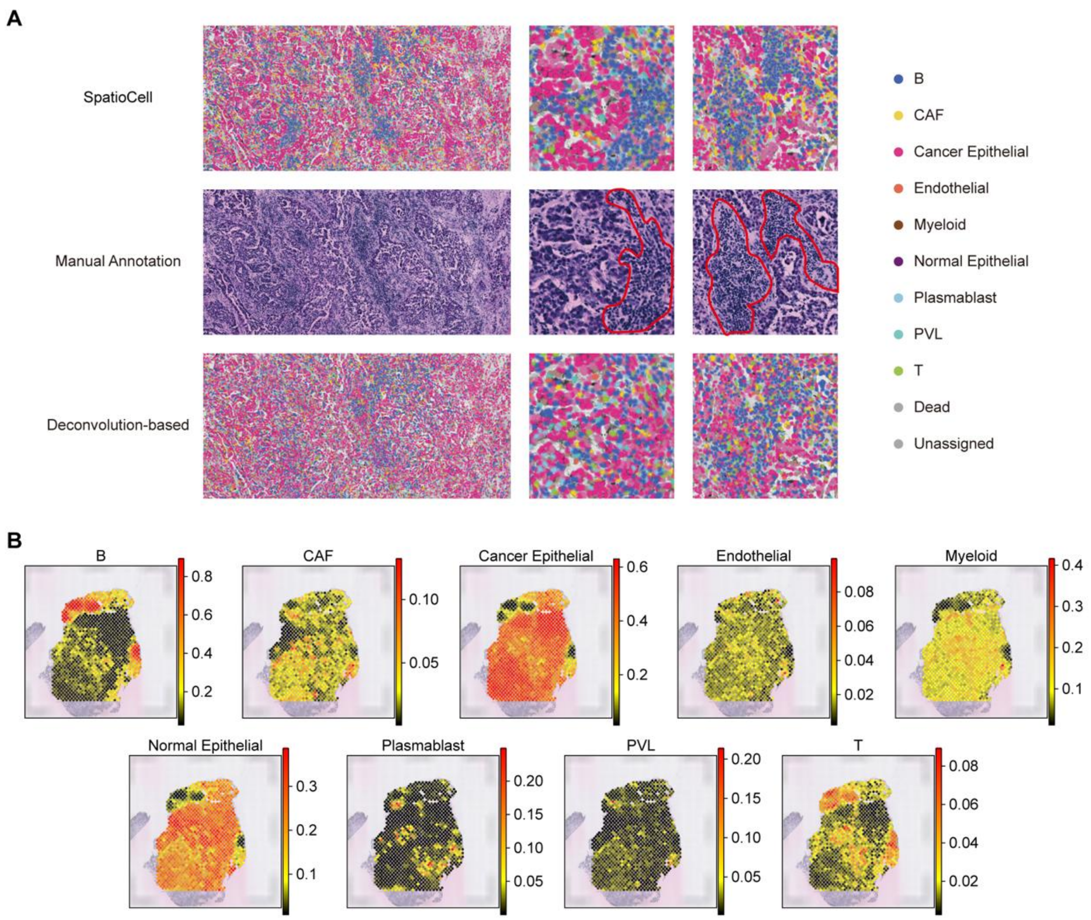


**Supplementary figure 7. Visualized cell type annotation results of SpatioCell in TNBC dataset.** A. Visualized cell type annotation results of SpatioCell in TNBC P30 data. Cells were first segmented in H&E image and then annotated using SpatioCell. B. Deconvolution results of TNBC P30 data using Cell2location, colored by cell abundance of each cell type.

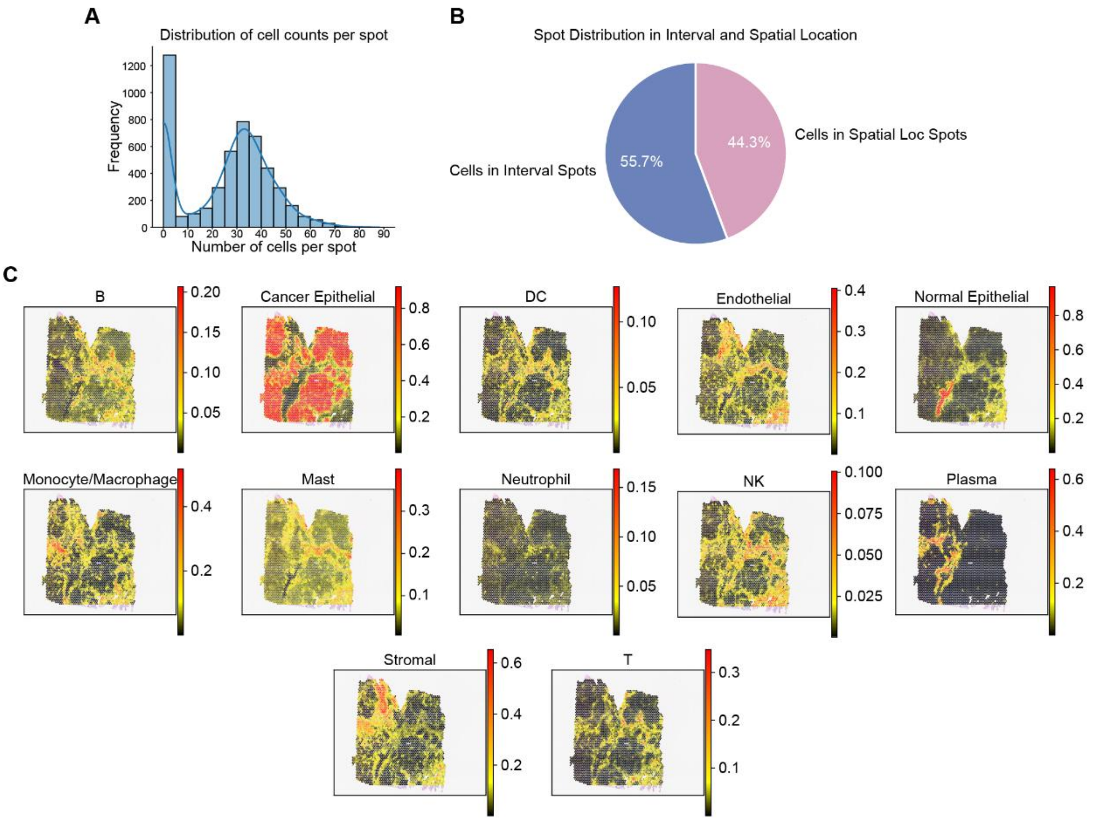


**Supplementary Figure 8. Cell distribution and deconvolution results of LUSC data.** A. Distribution of cell counts in captured regions of a lung squamous cell carcinoma (LUSC) sample sequenced with 55 μm spot size. B. Proportion of total cell counts between captured and inter-spot regions in the LUSC sample. C. Cell abundance of each cell type in lung cancer Visium dataset deconvoluted by Cell2location.
